# The nucleus activates mechano-responsiveness via FHOD-associated LINC complexes

**DOI:** 10.1101/2025.06.13.659598

**Authors:** Susumu Antoku, Leroy C. Joseph, Prachi Dabhade, B. Xiaokan Zhang, Chyuan-Sheng Lin, Thomas U. Schwartz, Jared Kushner, Barry M. Fine, John P. Morrow, Howard J. Worman, Gregg G. Gundersen

**Affiliations:** Department of Pathology & Cell Biology, Vagelos College of Physicians & Surgeons, Columbia University, New York, NY 10032; Department of Medicine, Vagelos College of Physicians & Surgeons, Columbia University, New Your, NY, 10032; Department of Biology, Massachusetts Institute of Technology, Cambridge, MA 02139; Herbert Irving Comprehensive Cancer Center, Vagelos College of Physicians & Surgeons, Columbia University, New York, NY 10032

## Abstract

The nucleus is the defining organelle of eukaryotic cells. It is usually considered a target organelle for cellular inputs. Here, we find that the nucleus is not simply a “passive” responder, but an active organelle directing the mechanical properties of the actin cytoskeleton it engages. Biochemically, interaction of FHOD formins with nesprin-2 of the nuclear LINC complex activates their actin bundling activity making them more potent than known bundlers like fascin or α-actinin. In cells, FHOD-associated LINC complexes enhance the mechanical resistance of nuclear-engaged actin cables in polarizing fibroblasts and sarcomeres in developing cardiomyocytes. Hypertrophic cardiomyopathy-associated variants of FHOD3 are defective in these processes. In mice, the *FHOD3* R637P disease-causing allele results in embryonic lethality when homozygous and in stress-induced cardiac hypertrophy when heterozygous. These results show that the nucleus actively directs its mechanical environment and that disruption of this capability in heart leads to cardiac hypertrophy.

## INTRODUCTION

The nucleus is the largest and stiffest organelle in eukaryotic cells (*1, 2*). In addition to defining the nuclear boundary and packing genomic DNA into a small space, the nuclear envelope is endowed with the ability to interact with cytoskeletal filaments through the linker of nucleoplasm and cytoskeleton (LINC) complex composed of inner nuclear membrane SUN proteins and outer nuclear membrane KASH (nesprin) proteins (*3, 4*). SUN domain proteins interact with the nuclear lamina and nesprins with cytoskeletal components. This allows cytoskeletal generated forces to position and move the nucleus, change its shape, transduce mechanical signals into the nucleoplasm, and move chromatin. The mechanical forces exerted on the nucleus for these activities range broadly, with very small pN forces capable of indenting the nucleus and initiating mechanotransduction to very large nN forces required to move it rapidly during the cell cycle or during cell polarization and migration (*2, 5, 6*).

The broad range of forces encountered by the nucleus requires distinct molecular assemblies to transmit them. Indeed, LINC complexes and associated cytoskeletal elements assemble into supermolecular arrays, such as TAN lines (*7–10*), the actin cap (*11*) and ALL lines (*12*), to transmit high forces from the actin cytoskeleton to the nucleus. Lower forces may be accommodated by individual LINC complexes (*13*). At the molecular level, SUN proteins and nesprins can form covalent disulfide bonds to resist higher forces and these are necessary for certain types of nuclear positioning (*14*).

Current models for nuclear coupling of cytoskeletal filaments assume that the attached filaments are not altered by their interaction with the LINC complex. However, recent studies on microtubule association with the nucleus have revealed that microtubule motors are activated by interaction with nesprins (*15, 16*). Despite the abundant evidence for actin filaments interacting with and exerting force on the nucleus, there is currently no evidence whether the actin filaments themselves are affected upon interaction with the nucleus. Other membrane-associated actin arrays, such as actin fibers associated with focal or cell-cell adhesions, there is strong evidence that the adhesion alters the array (*17–19*). Additionally, for many actin-nesprin interactions, the interaction is not simply mediated by the paired calponin homology domains of nesprin- 1/2, but also involves nesprin1/2-associated factors such as the formins FHOD1/3 (*8, 20*), fascin (*21*) and Mena (*22*). Each of these proteins have their own actin bundling and/or polymerization activity that could affect actin arrays.

The LINC complex and associated proteins such as the nuclear lamins and FHOD3 function in the transmission of forces in cells subjected to recurrent mechanical stress such as those in the heart. Mutations in *LMNA* encoding lamin A/C cause dilated cardiomyopathy (*23, 24*). SUN and nesprin variants have similarly been linked to cardiomyopathy (*25, 26*). Mutations in FHOD3 have also been linked to hypertrophic and dilated cardiomyopathy (*27–31*). Studies of cardiac sarcomere formation reveal a close association with the nucleus and in iPSC-derived cardiomyocytes, FHOD3 is required for the formation of sarcomeres (*32, 33*). Therefore, the mechanical coupling of actin to the nucleus via LINC complexes may play a particularly important role in sarcomere genesis and protecting the heart from disease.

Here, we show that nesprin interaction with the FHOD1/3 family of bundling formins profoundly affects their bundling activity and changes the mechanical behavior of actin filaments engaged by the nucleus. In cardiomyocytes, this leads to defects in sarcomere assembly. We further show that in mice homozygosity for a FHOD3 variant that interferes with sarcomere assembly leads to embryonic lethality at the time of heart development whereas heterozygosity leads to stress-induced cardiac hypertrophy.

## RESULTS

### Nesprin2-G spectrin repeat (SR) 11-13 activates the F-actin bundling activity of FHOD1/3

FHOD1’s amino terminus interacts with SR 11-12 of nesprin-2G forming a high affinity complex with two contact sites that are characterized by salt bridges between the two proteins (*34*). This interaction is required for nesprin-2-and actin-dependent movement of the nucleus in fibroblasts polarizing for migration (*34*). A more recent AlphaFold 3 structure shows that SR11-12 are part of a continuous structural element encompassing SR11-13 of nesprin-2G (denoted SR11-13) (fig. S1A). The FHOD1-SR11-13 complex structure maintains the original interacting site residues and does not reveal additional contact sites. However, a thermal stability assay revealed that the addition of SR13 to SR11-12 stabilizes the extended SR11-13 spectrin module (fig. S1B). Biolayer interferometry binding assays confirm that SR11-13 interacts with FHOD1 with higher affinity compared to SR11-12 (K_D_ = 92 nM vs 472 nM), apparently due to a lower off rate (fig. S1C). GST-pulldown assays also confirmed the higher affinity of nesprin-2 SR11-13 to FHOD1 than nesprin-2 SR11-12 (fig. S2A).

As SR11-13 is more likely to represent the structure of the native protein, we used it to test whether nesprin-2G interaction with FHOD1 affected its actin filament bundling and polymerization activities (*20, 35, 36*). To measure bundling, we used a low-speed centrifugation assay that separates actin bundles from individual actin filaments (F-actin). We defined actin bundling efficiency as the concentration of protein that bundled 50 % of 2 uM F-actin. FHOD1 wild-type (WT) alone did not exhibit detectable F-actin bundling activity at 500 nM (Fig. 1A and B), consistent with its autoinhibited state. In contrast, when SR11-13 was added there was a dramatic gain in F-actin bundling activity; bundling efficiency was now 50 nM FHOD1 (Fig. 1A and B).

**Figure 1.**
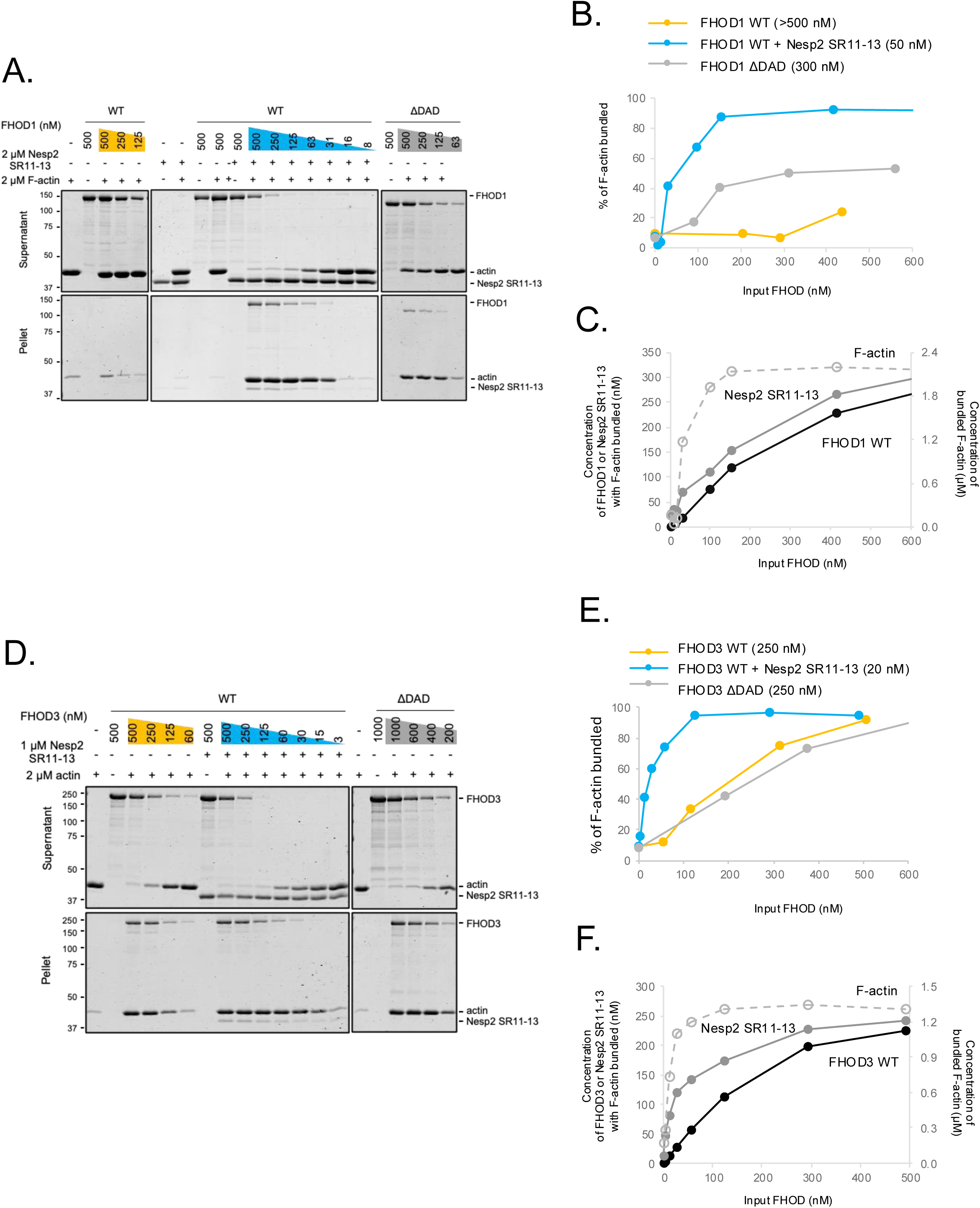
Nesprin-2G SR11-13 stimulates the F-actin bundling activity of FHOD1/3. **A** and **D.** Coomassie stained SDS polyacrylamide gel from *In vitro* F-actin bundling assay in which actin bundles are pelleted, but not individual filaments. The indicated purified proteins were incubated with F-actin and F-actin bundles were pelleted at 16,000 x *g*. Supernatant and pellet fractions are shown for the indicated input proteins. Molecular mass markers in kDa are indicated on the left. Nesp2 SR11-13 = nesprin-2 SR11-13. **B** and **E**. Concentration dependence of F-actin bundling efficiency by FHOD1 (B) and FHOD3 (E). Bundled F-actin and FHOD were quantified from gels as in A and D. The number in the parenthesis in the sample legend indicates the F-actin bundling efficiency, defined as the concentration of FHOD required for bundling 50 % of the F-actin. **C** and **F.** Stoichiometry of FHOD and nesprin-2 SR11-13 in the bundled F-actin fraction. The amounts of actin, FHOD, and nesprin-2 SR11-13 were quantified from gels as in A and D.

This represents a >10-fold increase in bundling efficiency over FHOD1 WT and is even 6-fold higher that that observed with the constitutively active form of FHOD1 in which the autoregulatory DAD is removed (FHOD1 ΔDAD) (Fig. 1A and B). Similar to FHOD1, FHOD3 binds to nesprin-2 SR11-13 (fig. S2A). SR11-13 also stimulated the bundling activity of FHOD3 (to 20 nM), although FHOD3 had an intrinsic bundling efficiency (250 nM) (Fig. 1D and E). Consistent with the biochemical data, FHOD1 WT plus SR11-13 generated thicker actin bundles than FHOD1 WT or FHOD1 ΔDAD by a TIRF microscopy assay (fig. S2B).

The SR11-13 stimulated FHOD1 F-actin bundling required interaction between the two, as mutants preventing interaction blocked stimulation of bundling and SR11-13 alone did not bundle (fig. S2C). SR11-13 did not stimulate the bundling activity of the formin mDia1, and SR51-54 of nesprin-2G, which interacts with a number of other proteins (*37, 38*), did not stimulate the bundling activity of FHOD1 showing the specificity of the bundling stimulation (fig. S3A and B).

Consistent with the importance of the triplet SR11-13, none of the individual SRs (SR11, SR12, SR13) or doublet SRs (SR11-12, 12-13) activated FHOD1 bundling activity (fig. S3C). The SR11-13 fragment was detected in the bundled F-actin fraction, but only when FHOD1 was also present, indicating that SR11-13 did not bind to F-actin itself. Supporting this interpretation, the stoichiometry of SR11-13 to FHOD1 WT and FHOD3 WT in the bundled F-actin fraction was 1:1.18 +/- 0.05 (SEM) and 1:1.3 +/- 0.11 (SEM), respectively (Fig. 1A, C, D, and F).

To gage relative efficiency of SR11-13-stimulated FHOD1/3 bundling of F-actin, we compared its bundling activity to that of well-known F-actin bundlers, fascin1 and α-actinin 1. The bundling activity of SR11-13 stimulated FHOD1 was 2-2.5-fold and FHOD3 5-6.3 fold higher than that of fascin1 or α-actinin1 (fig. S4A and B). Thus, SR11-13 stimulates the F-actin bundling activity of FHOD1/3 to levels well above those of known actin bundling proteins.

### Nesprin2-G SR11-13 does not stimulate the F-actin polymerizing activity of FHOD1/3

Unlike its effect on bundling activity, SR11-13 did not stimulate FHOD1/3’s actin polymerization activities as measured with the pyrene actin polymerization assay (fig. S4C and D). As reported previously (*20, 35*), the constitutively active FHOD1 ΔDAD stimulated actin polymerization, but to a considerably lower level compared with that of mDia1 ΔDAD (fig. S4C and D). The conserved Ile residue in the FH2 domain that is essential for actin polymerization by other formins was also required for FHOD1 stimulated actin polymerization (fig. S4C and D). There was little detectable difference in FHOD1 actin polymerization activity using either muscle or non-muscle actin. Overall, these results show that nesprin-2G SR11-13 selectively activates FHOD1’s actin bundling activity without activating its actin polymerization activity.

### Nesprin-2G SR11-13 activates a cryptic actin binding site in FHOD1/3

As far as we are aware, the stimulation of FHODs’ F-actin bundling activity by interacting with another protein is unique among F-actin bundling proteins. We pursued the mechanism for SR11-13 stimulated FHOD1 bundling. Bundling by FHOD1 requires its FH2 domain, although not its barbed end binding capability, and a second site in the intrinsically disordered region (IDR) near its amino terminus (*20, 36, 39*). We tested whether the interaction between SR11-13 and the amino terminus might “activate” a cryptic actin binding site in the IDR using a high-speed actin pelleting assay. As expected, SR11-13 stimulated the actin binding of full length FHOD1, and SR11-13 alone did not exhibit F-actin binding (Fig. 2A-C). Neither of the FHOD1 amino terminal constructs, SRBD-DID or SRBD-DID-IDR, bound actin alone (Fig 2D and E). Addition of SR11-13 stimulated the F-actin binding of SRBD-DID-IDR, but not that of SRBD-DID (Fig 2 D, and E). In contrast to FHOD1, FHOD3 SRBD-DID-IDR bound F-actin by itself, although addition of SR11-13 further stimulated its F-actin binding (Fig. 2D and E).

**Figure 2.**
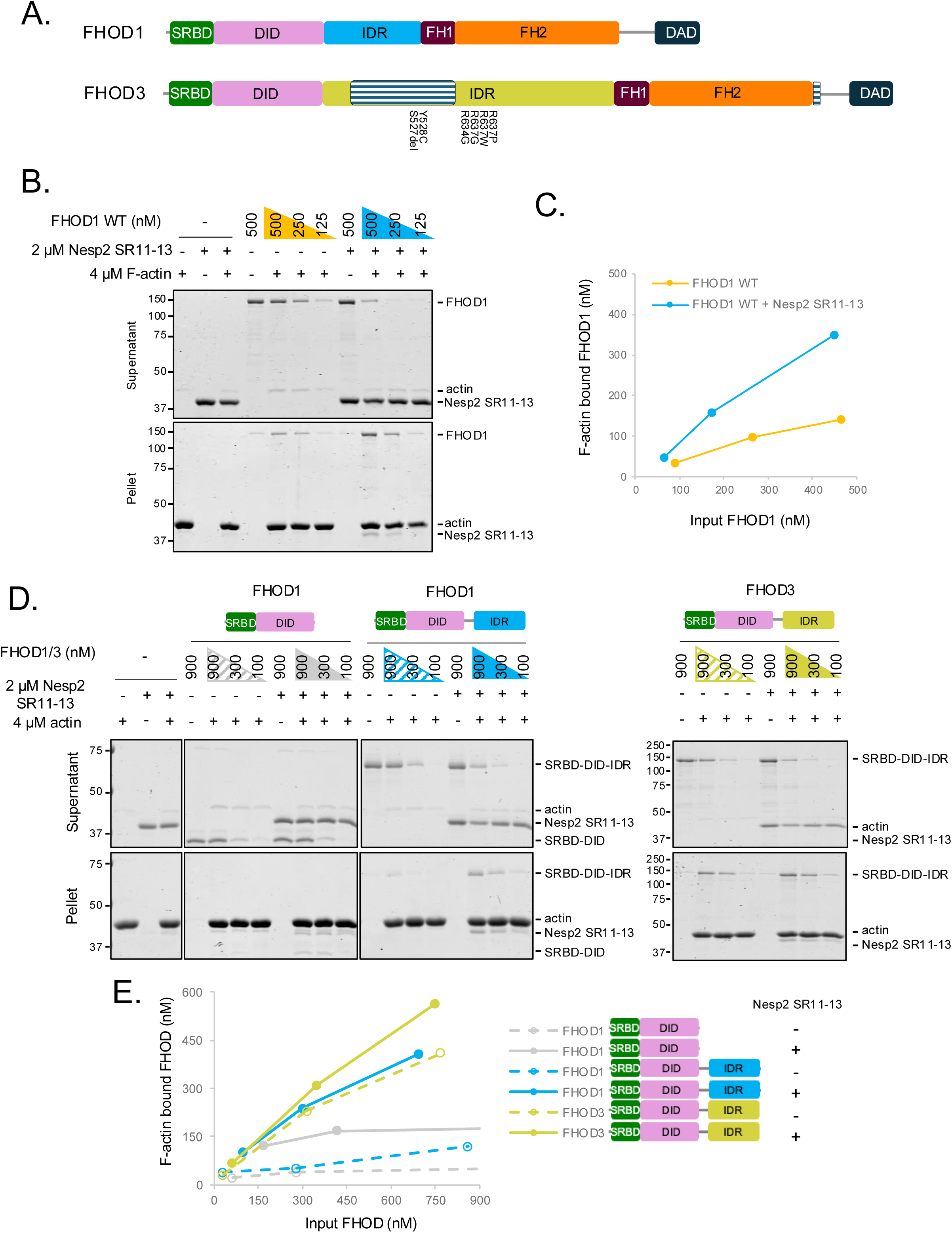
Nesprin-2G SR11-13 promotes F-actin binding activity of FHOD1/3 SRBD-DID-IDR. **A.** Diagram of FHOD1/3. SRBD, DID, IDR, FH1/2, and DID are SR binding domain, diaphanous inhibitory domain, intrinsically, disordered region, Formin Homology domain 1 and 2, and diaphanous auto-regulatory domain, respectively. Cardiac specific insert region of FHOD3 is shown in the striped box. The amino acid change below the FHOD3 diagram shows the site of hypertrophic cardiomyopathy disease variants examined in this study. **B** and **D**. F-actin binding by full-length FHOD1, and amino terminal constructs from FHOD1 (SRBD-DID, and SRBD-DID-IDR) and FHOD3 (SRBD-DID-IDR) proteins in the presence or absence of nesprin-2 SR11-13 examined by a high-speed (100,000 x *g*) F-actin co-sedimentation assay. Coomassie blue-stained SDS-polyacrylamide gels of proteins in the supernatant and pellet are shown. Concentrations of proteins are as indicated. Molecular mass standards in kDa are indicated at the left. **C** and **E**. Concentration dependence of full-length FHOD1 (C) and amino terminal constructs of FHOD1/3 (E) binding to F-actin with and without added nesprin-2G SR11-13. Protein quantification was by scanning gels such as that in B, D.

Given the lack of SR-11-13 binding to F-actin (see Fig. 2B and D), these results strongly suggest that nesprin-SR11-13 interaction with FHOD1 induces a cryptic actin binding activity in the amino terminus of FHOD1 and that this enhanced binding likely involves the IDR. Consistent with this, ERK1/2 phosphorylation of the IDR and phospho-mimetic mutants of the IDR do not bundle F-actin (*20*).

### FHOD1 reinforces F-actin cables engaged by the nucleus during nuclear movement

The stimulation of actin bundling of FHOD1 by SR11-13, suggests that FHOD1 may enhance mechanical resilience of actin filaments connected to the nucleus by nesprin-2G-FHOD1 complexes. To test this idea, we examined dorsal actin cables that engage the nucleus via nesprin-2G and FHOD1 to move it rearward in fibroblasts polarizing for migration (*8–10, 20*). We monitored strain on actin cables expressing EGFP-tagged zyxin-LIM domain (EGFP-zyxin-LIM), which selectively decorates strained actin bundles (*40–43*), and Lifeact-mScarlet to label F-actin. Lysophosphatidic acid (LPA) stimulation of serum-starved NIH3T3 fibroblasts triggered formation of dorsal actin cables in the leading lamella and their flow rearward as previously described (*9, 44*) (Supplemental Movie 1A). EGFP-zyxin-LIM selectively decorated dorsal actin cables that were localized above the nucleus, with lower frequency of labeling elsewhere in the cell (Fig. 3A and B). Sites of individual zyxin-LIM patches correlated with thinner regions of the actin cables, consistent with their specificity for strained actin cables (Fig. 3A, arrows). Strain detected by EGFP-zyxin-LIM temporally correlated with nuclear movement, peaking at 60 min and then subsiding until the nucleus stopped moving (Fig. 3B; Supplemental Movie 1A and 1B). This spatial-temporal correlation strongly suggests that the moving nucleus is responsible for the appearance of strain in these actin cables.

**Figure 3.**
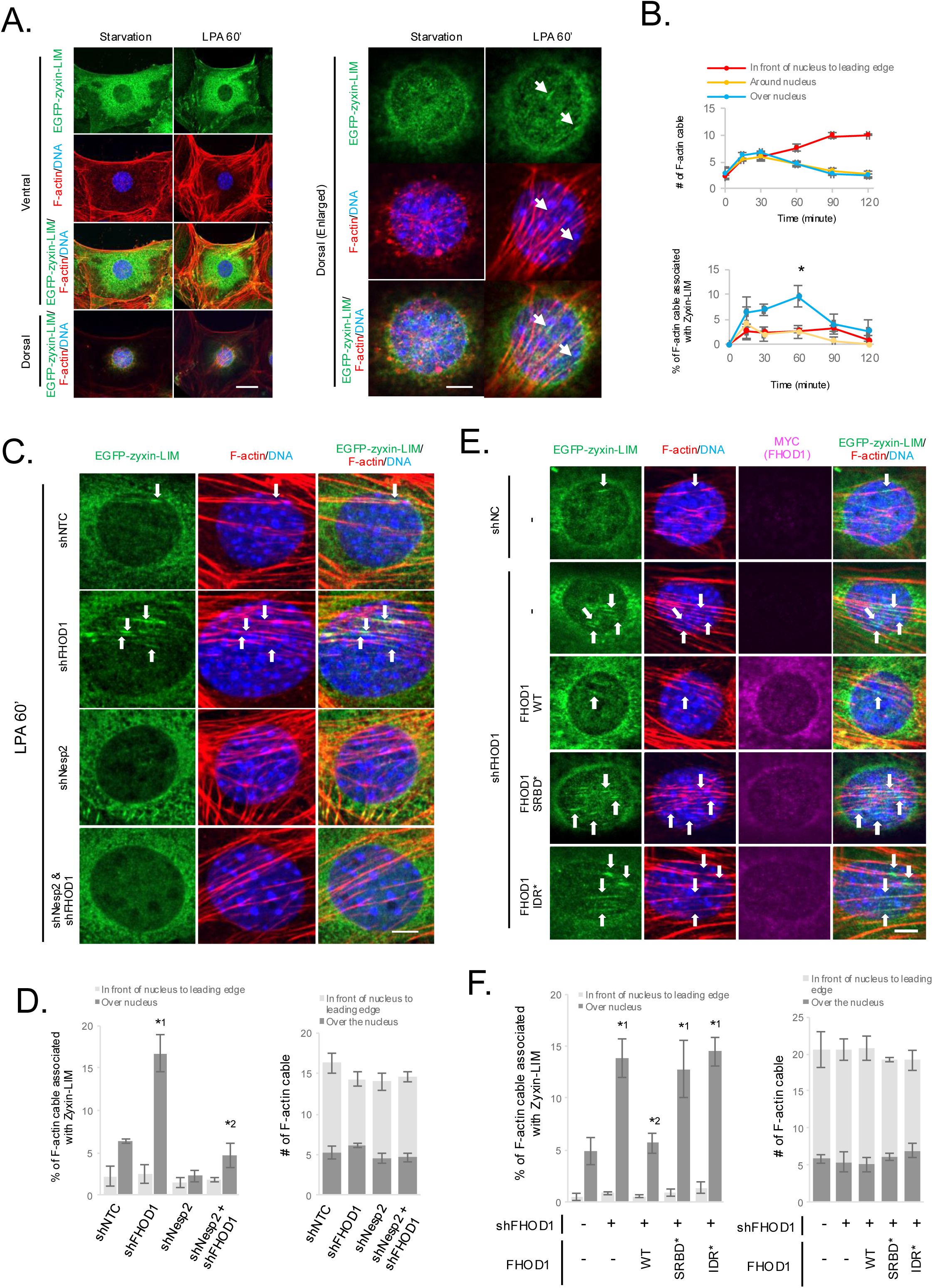
FHOD1 is required to prevent strain in actin cables over nucleus. **A**. Images of serum-starved NIH3T3 fibroblasts immunofluorescently stained for the indicated proteins after expressing doxycycline inducible EGFP-zyxin-LIM and treatment with or without LPA for 60 min. Bar, 20 μm. Higher magnification views of the nucleus are shown in the panels on the right. Bar 5 μm. White arrows, zones of increased EGFP-zyxin-LIM decoration coupled with thinned F-actin cables indicating strained F-actin. **B**. The frequency of strained actin cables (EGFP-zyxin-LIM decoration and thinned actin cable) filament and strained F-actin at the different subcellular locations after LPA stimulation for the indicated times. The mean and SEM is shown from at least 3 independent experiments, n = 15 to 25 cells/experiment. * indicates p < 0.05 compared to the strained F-actin from other two areas at a given time. **C-F**. (**C** and **E**) Immunofluorescence images of NIH3T3 fibroblasts knocked down for indicated proteins after expressing doxycycline inducible EGFP-zyxin-LIM. Starved cells were stimulated with LPA for 1 h. White arrows, increased EGFP-zyxin-LIM decoration coupled with thinned F-actin filament indicating strained F-actin. Bar, 5 μm. (**D** and **F**) The frequency of strained F-actin at the different subcellular locations in cell treated with the indicated shRNAs (shNTC and shNC are a noncoding control). In (E and F), FHOD1 knockdown NIH3T3 fibroblasts stably expressing the indicated FHOD1 rescue constructs. The mean and SEM is shown from at least 3 independent experiments, n = 20 to 25 cells/experiment. In (**D**), *^1^ indicates p < 0.05 compared to nuclear position of shNTC NIH3T3 fibroblast. *^2^ indicates p < 0.05 compared to nuclear position of shFHOD1 NIH3T3 fibroblast and p > 0.05 compared to nuclear position of shNTC NIH3T3 fibroblast. In (**F**), *^1^ indicates p < 0.05 compared to nuclear position of shNC NIH3T3 fibroblast. *^2^ indicates p < 0.05 compared to nuclear position of shFHOD1 NIH3T3 fibroblast and p > 0.05 compared to nuclear position of shNC NIH3T3 fibroblast.

Knockdown of FHOD1 increased the frequency of strained actin cables detected by EGFP-zyxin-LIM over the nucleus without affecting actin cables distal to the nucleus or the total number of actin cables (Fig. 3C and D and fig. S5A). When the connection between the nucleus and the dorsal actin cables was disrupted by knocking down nesprin-2, strain in the actin cables over nucleus was reduced compared to controls, although the change was not statistical significance (Fig. 3C and D and fig. S5A). Furthermore, when both nesprin-2 and FHOD1 were knocked down, the frequency of strained F-actin over the nucleus decreased. These data indicate that the strain on the actin cables over the nucleus develops because of their attachment to the nucleus.

To determine whether the F-actin strain resistance was due to specific activities of FHOD1, we monitored strain in FHOD1-depleted cells re-expressing FHOD1 point mutants (Fig. 3E and F and fig. S5B). Re-expression of FHOD1 WT in knocked down cells significantly reduced strain in nuclear actin cables compared the knockdown cells. Re-expression of FHOD1 SRBD*, a variant that cannot bind to nesprin-2 (*34*), or FHOD1 IDR*, a variant that cannot bundle F-actin (*20*), failed to reduce the elevated strain on nuclear actin cables in the FHOD1 knockdown cells.

To test whether enhancing the repair of strained actin cables was sufficient to restore nuclear movement in FHOD1 depleted fibroblasts, we overexpressed full length zyxin in FHOD1 knockdown cells. zyxin over-expression rescued the nuclear movement defect in FHOD1 knockdown cells (fig. S5C-E). Taken together, these data show that the essential function of FHOD1 in actin- and LINC complex-dependent nuclear movement is to reinforce actin cables against excessive strain caused by moving the large nucleus.

### FHOD3-nesprin interaction is required for proper sarcomere formation in cardiomyocytes

Like FHOD1, FHOD3 bundles F-actin, binds nesprin-2G SR11-13 and exhibits enhanced bundling activity in the presence of nesprin-2 SR11-13 (see Fig. 1D). Unlike the widespread expression of FHOD1, FHOD3 is primarily expressed in cardiomyocytes where it is necessary to generate cardiomyocyte sarcomeres (*33, 45, 46*). As a result, FHOD3 knockout mice are early embryonic lethal (*46*). To determine whether these phenotypes are related, we examined whether FHOD3 interaction with the nucleus was critical for sarcomere formation in human induced pluripotent stem cell-derived human cardiomyocytes (iPSC-CMs).

FHOD3 knockdown in iPSC-derived CMs strongly disrupted formation of sarcomeres and impaired contraction (Fig. 4A, fig. S6A and B and Supplemental Movie 2A). Expression of FHOD3 WT-EGFP rescued sarcomere formation as previously reported (Fig. 4A) (*45*). FHOD3-EGFP localized to the sarcomere and to the dorsal aspect of the nuclear surface (Fig. 4A). Expression of a nesprin-2-binding deficient mutant of FHOD3 (FHOD3 SRBM*-EGFP) failed to rescue sarcomere formation and to localize to the nucleus (Fig. 4A).

**Figure 4.**
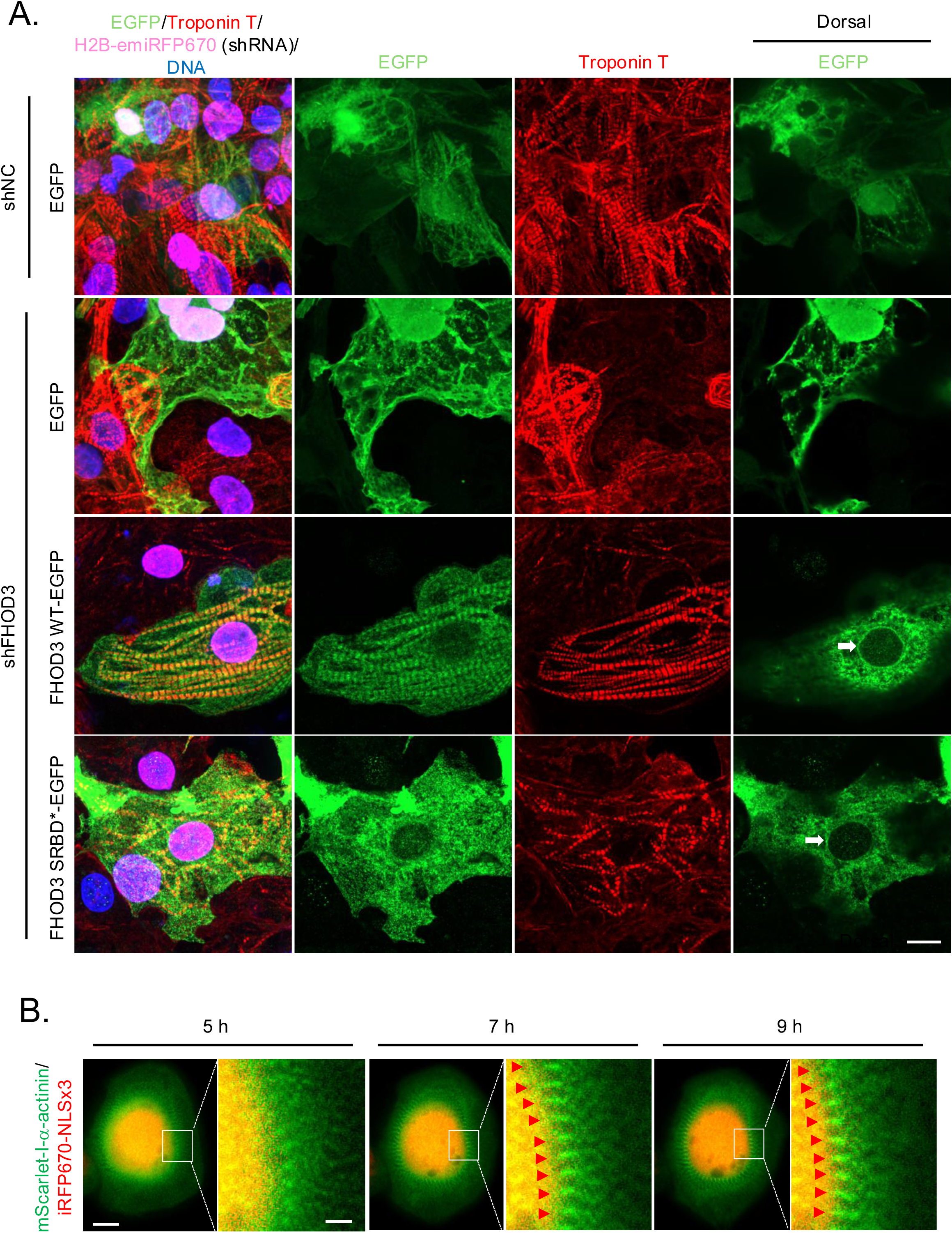
FHOD3-nesprin interaction is required for formation of organized sarcomeres in cardiomyocytes. **A**. Immunofluorescence images of iPSC-CM expressing the indicated proteins and shRNA and stained for the indicated proteins and DNA (DAPI). iPSC-CM were infected with lentiviruses expressing the indicated shRNA and H2B-emiRFP670 and one carrying doxycycline-inducible indicated EGFP or FHOD3-EGFP. The cells were replated on glass-bottom dish and the expression of EGFP or FHOD3-EGFP was induced by the addition of 100 ng/ml doxycycline for 4 d. White arrows, FHOD3 localization on the dorsal surface of the nuclear envelope. Bar, 10 μm. **B**. Panels from a movie of sarcomere formation after replating iPSC-CM cells. iPSC-CM were transfected with plasmids expressing Lifeact-EGFP, mScarlet-I-α-actinin, and iRFP670-NLSx3. Cells were detached, replated and then imaged by epifluorescence microscopy. The boxes denote the region shown at higher magnification at each time point. Red arrow heads, sites of sarcomere formation in close proximity to the nucleus. Bar, 5 μm. Bar in high magnification image, 1 μm.

These results suggest that FHOD3 must interact with nesprin-2 SR11-13 to contribute to sarcomere formation. Despite FHOD3 localizing to the nucleus, sarcomeres did not when examined 4 d after induction of FHOD3 expression to rescue the phenotype. To look at sarcomere formation dynamically, we used a replating assay with iPSC-CMs in which sarcomeres that are lost upon detachment, rapidly form after replating (*33, 47*). At early times after replating (5 h), sarcomeres are detected near the nucleus with sarcomere z-discs projecting horizontally to the nuclear surface (Fig. 4B, Supplemental Movie 2B). Over time, the sarcomeres become increasingly apposed to the nuclear surface. These data support the idea that sarcomeres form in association with the nucleus.

### Hypertrophic cardiomyopathy disease-causing variants of FHOD3 are defective in assembling mechanically engaged nuclear actin arrays in fibroblasts and cardiomyocytes

Mutations in *FHOD3* cause inherited hypertrophic cardiomyopathy (*28, 30*). The mutations cluster around the IDR of FHOD3 and include recurrent *FHOD3* mutations in at least three unrelated families (Fig. 2A). Although the genetics strongly support these mutations as disease causing, there are no studies of whether they affect FHOD3 function. To explore this, we first expressed a set of the FHOD3 variants in NIH3T3 fibroblasts depleted of FHOD1. Whereas FHOD3 WT rescued the nuclear movement in FHOD1-depleted cells, most of the disease variants failed to do so (Fig. 5A and B). The variants that failed to rescue were missense mutations in the IDR. We checked one of these, FHOD3 R637P to see whether it would rescue sarcomere formation in FHOD3 knockdown iPSC-CMs. FHOD3 R637P-EGFP localized to the nuclear envelope in iPSC-CM, however, it failed to rescue the formation of organized sarcomeres (Fig. 5C).

**Figure 5.**
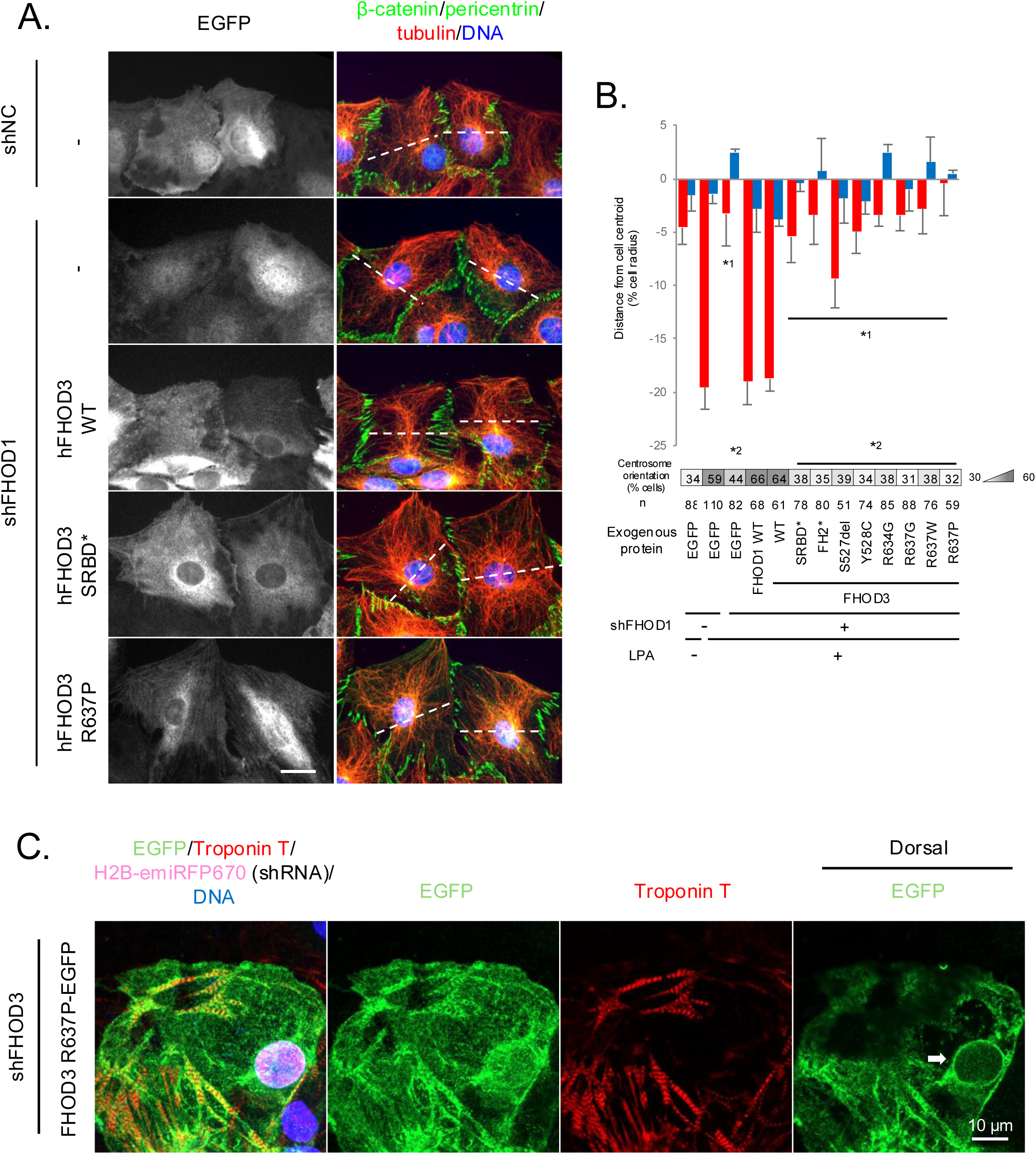
Disease causing mutations in FHOD3 impairs FHOD3 activity required for nuclear movement in fibroblasts and sarcomere organization in iPSC-CM. **A**. Immunofluorescence of the indicated proteins in a wounded monolayer of serum-starved NIH3T3 fibroblasts expressing doxycycline inducible EGFP or EGFP-tagged FHOD and scrambled shRNA or shRNA targeting FHOD1 after LPA stimulation for 2 h. The images were taken by epifluorescence microscopy. Dashed lines show the middle line of cell. Bar, 20 μm. **B**. Histogram of centrosome and nuclear positions relative to the cell centroid (defined as “0”; negative values toward the cell rear). Centrosome orientation (heat map below the histogram, values mean % of cells) toward the front of the cell for cells treated as in **A**. Values are means ± SEM; n, cells examined. *^1^ and *^2^ indicates p < 0.05 compared to nuclear position and centrosome orientation, respectively, of shFHOD1 NIH3T3 fibroblast with EGFP expression. **C**. Immunofluorescence images of iPSC-CM expressing FHOD3 R637P-EGFP and shRNA targeting FHOD3. iPSC-CM were infected with lentiviruses expressing shRNA targeting FHOD3 and H2B-emiRFP670 and one carrying doxycycline-inducible FHOD3 R637P-EGFP. The cells were replated on glass-bottom dish and the expression of FHOD3 R637P-EGFP was induced by doxycycline for 4 d. White arrow, localization of FHOD3 R637P-EGFP to the dorsal surface of nuclear envelope. Bar, 10 μm.

### Fhod3^R611P/+^ mice exhibit cardiac hypertrophy

Autosomal dominant mutations in *FHOD3* have been linked to hypertrophic cardiomyopathy in humans (*28, 29*). We therefore generated *Fhod3* knock-in mice with a R611P (corresponding to R637P in humans) mutation (fig. S7A). Knock-in of the *Fhod3* mutation was confirmed by PCR (fig. S7B) and DNA sequencing and showed heterozygous insertion (fig. S7C). *Fhod3^+/+^*and *Fhod3*^R611P/+^ mice were born at the expected Mendelian ratios but no *Fhod^R611P/R611P^* mice were born (fig. S8A).

*Fhod3^R611P/R611P^* embryos were only detected up to embryonic day 11.5 (fig. S8B). This suggested that the FHOD3 R611P variant was null or dominant negative for cardiac development. Immunoblotting showed that the expression of FHOD3 was the same in *Fhod3*^+/+^ and *Fhod3*^R611P/+^ mice, suggesting the FHOD R611P mutation did not alter protein stability (fig. S8C). Growth of *Fhod3^R611P/+^*mice was similar to *Fhod3*^+/+^ mice when followed for 10 months (fig. S8D). *Fhod3^R611P/+^* mice had normal left ventricular diameters and function at 3-6 months as assessed by echocardiography (fig. S8E). At 3 and 6 months of age, there were also no differences in measured heart to body mass ratio (fig. S8F) or left ventricular mass to body mass as assessed by echocardiography (fig. S8G).

As there were no detectable differences between *Fhod3^R611P/+^*and *Fhod3*^+/+^ mice, we hypothesized that stressing the heart by infusion of angiotensin II would induce more severe cardiac hypertrophy in the mutant mice. Angiotensin II infusion increased gross heart size in both *Fhod3*^+/+^ and *Fhod3^R611P/+^* mice compared to sham-treated controls (Fig. 6A). Compared to angiotensin II-treated *Fhod3*^+/+^ mice, angiotensin II-treated *Fhod3^R611P/+^*mice had an increase in measured heart to body mass ratio (Fig. 6B) and left ventricular mass to body mass as assessed by echocardiography (Fig. 6C and D). Echocardiography showed increased left ventricular anterior wall thickness in angiotensin II-treated compared to sham-treated *Fhod3^R611P/+^* mice but no increased was seen in angiotensin II-treated *Fhod3*^+/+^ mice (Fig. 6E and F). Consistent with these morphological findings, there were significant increases in the expression of hypertrophic markers *Nppa* and *Nppb* encoding atrial natriuretic peptide and brain natriuretic peptide, respectively (Fig. 6G and H). Microscopic examination of left ventricle sections stained with Masson’s trichrome showed fibrosis in hearts of both *Fhod3*^+/+^ and *Fhod3^R611P/+^* mice (Fig. 6I). However, the fibrotic area was significantly greater in left ventricles of *Fhod3^R611P/+^* mice than *Fhod3*^+/+^ mice after treatment with angiotensin II (Fig. 6J). Consistent with the histopathological findings, expression of mRNAs encoding collagen I, and collagen III were significantly increased in left ventricular tissue from *Fhod3^R611P/+^* mice compared to *Fhod3*^+/+^ mice after angiotensin II treatment (Fig 6K and L). However, left ventricular fractional shortening and ejection volume were unaffected by angiotensin II-treatment of *Fhod3*^+/+^ and *Fhod3^R611P/+^* mice (fig. S9A and B). These data show that the *Fhod3^R611P/+^* mice are prone to cardiac hypertrophy upon stress.

**Figure 6.**
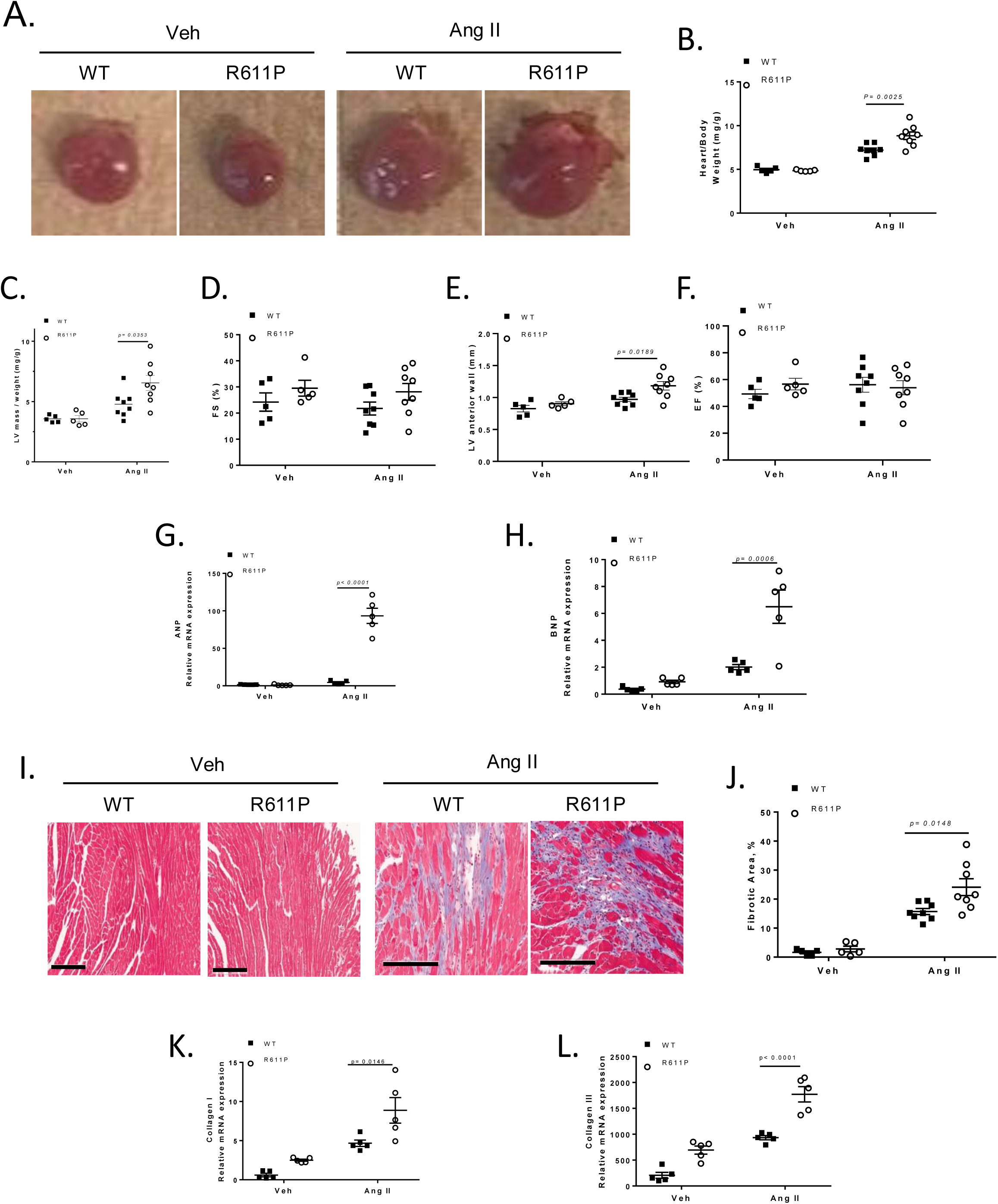
Mutation in *Fhod3* exacerbates Ang II-induced Cardiac hypertrophy and fibrosis. Mice was subjected to subcutaneous infusion of 2.5 mg/kg/day Ang II by Osmotic pump for 28 days. **A**. Representative images of heart size. **B**. Quantification of heart weight normalized to body weight (n=5-8). **C**. Left ventricular mass normalize to body weight (n=5-8). **D**. Fractional shortening, left ventricle volume. **E**. Left ventricle anterior wall thickness (n=5-8). **F**. Left ventricle ejection fraction. **G** and **H**. Atrial natriuretic peptide RNA expression the ventricles. **I**. Representative images of Mason’s trichrome stained heart sections (scale bar: 200 μm). **J**. Quantification of myocardial collagen deposits (n-5-8). Data are presented as mean ± SEM, with n representing number of animals. p < 0.05 was considered statistically significant. **I** and **J**. Collagen RNA expression in the ventricles.

## DISCUSSION

Giant nesprins, such as nesprin-1G and nesprin-2G with their many SRs, have previously been considered as scaffolding proteins that endow the nucleus with the ability to bind to cytoskeletal and signaling proteins (*3*). However, our current results show that nesprin-2G’s SR11-13 directly activate the actin bundling activity of FHOD1 and FHOD3, indicating that the nucleus is an active participant in regulating the mechano-responsiveness of the actin cytoskeleton in its vicinity. The lack of this activity has consequences in cells and in mice as fibroblasts fail to move their nucleus, cardiomyocytes fail to develop sarcomeres and mice develop cardiac hypertrophy.

Coupled with previous data on nesprins activating microtubule motors (*15, 16*), these results show that the nucleus should be considered a more active organelle than previously thought. In the case of FHOD1/3, our data support a biomechanical effect on the actin cytoskeleton rather than one that affects actin polymerization dynamics. However, FHOD1/3 possess actin polymerizing activity (*20, 35*) and this activity might be important if FHOD1/3 were to also repair strained actin filaments, as suggested by the ability of overexpressed zyxin, an actin repair protein, to rescue nuclear movement in FHOD1-depleted fibroblasts.

Our results extend a study in developing Drosophila muscle implicating the nucleus and the LINC complex in sarcomere formation (*32*). We find that FHOD3 is localized to the nucleus during sarcomere formation and that disrupting this nuclear localization by mutating FHOD3’s nesprin-2 binding site, prevents normal sarcomere formation. We also observe the close association of the nucleus with forming sarcomeres, a result that was previously observed, but not followed up (*33, 47*). We hypothesize that developing sarcomeres engage the nucleus as a way to mechanically resist the forces generated by the developing sarcomeres so that sarcomeres can mature. In the absence of this association, force generation by the nascent sarcomeres may rip them apart generating the fragmented sarcomeres we detect in cells expressing FHOD3 that cannot associate with the nucleus (Fig. 4A). As sarcomeres in mature cardiomyocytes are not nuclear associated, we also hypothesize that there is a developmental step in which they are released from the nucleus.

Our results strongly support the dysfunction of *FHOD3* mutations that were previously shown genetically to cause hypertrophic cardiomyopathy (*28, 30, 31*). Our results on a mouse model show that the *Fhod3* R611P mutation, corresponding to the human *FHOD3* R637P hypertrophic cardiomyopathy-causing mutation, renders mice susceptible to left ventricular hypertrophy. This variant likely leads to loss of FHOD3 function, as the *Fhod^R611P/R611P^* mice die during embryogenesis at the same embryonic age reported for *Fhod^-/-^* mice (*46*). More work is needed to understand the precise mechanism by which the FHOD3 mutants disrupts function in adult hearts. Based on the clustering of many of the mutations in the IDR, the inability of mutations in the IDR to rescue nuclear movement in fibroblasts and the evidence for a cryptic actin binding site in the IDR, we suggest that the bundling activity required to mechanically reinforce actin filaments is the key activity. In developing cardiomyocytes, this activity would be essential for the formation of sarcomeres whereas in adults, the activity may be important to maintain or repair sarcomeric actin cables. Consistent with this, conditional knockout of FHOD3 in the heart, results in defects in cardiac function including hypertrophy (*48*).

Our study describes the first function for a formin’s actin bundling activity *in vivo*. Nonetheless, there is abundant evidence that other formin family members, usually considered regulators of actin nucleation and polymerization, also have actin bundling activity (*49–58*). This includes many mammalian formins but also those in other organisms. Activities of FHOD1/3 we document in the current study have been described earlier as individual properties of formin family members from phylogenetically divergent species. For example, ForC in *Dictyostelium* lacks the FH1 domain and actin filament polymerizing activity and instead is a dedicated actin bundling protein (*52*). The plant formin AtFH8 bundles actin filaments and is a dedicated outer nuclear membrane protein due to its encoded amino terminal transmembrane domain (*58*). These properties suggest a re-examination of the formin family may reveal conservation of features of this class of proteins other than its actin polymerizing activity.

## MATERIALS AND METHODS

### Cell culture

NIH3T3 fibroblasts were maintained in Dulbecco’s Modified Eagle Medium (DMEM; Corning Inc.) containing 10 mM HEPES pH 7.4, 10% (v/v) bovine calf serum (GE Health Life Science), 1X Penicillin Streptomycin (ThermoFisher). 293T cells were maintained in DMEM containing 10 % (v/v) fetal bovine serum (Gemini Bio-Products) and 1X Penicillin Streptomycin. iCell® Cardiomyocytes2 (FUJI FILM Cellular Dynamics Inc.) were cultured according to the manufacture’s protocol with supplied reagents.

Human induced pluripotent stem cells (hiPSC) were obtained through Material Transfer Agreements from Bruce Conklin, Gladstone Institute (WTC cell line). hiPSCs were expanded on growth factor reduced Matrigel-coated plates (Corning) in mTeSR Plus medium (STEMCELL Technologies) containing mTeSR supplement (STEMCELL Technologies), 1X Penicillin Streptomycin. Cell culture medium was changed every other day, and cells passaged upon reaching 70% confluence. During the first 24 h after passaging, the culture media was supplemented with 5 μM Y-27632 dihydrochloride (Tocris).

Cardiomyocytes (CM) were differentiated from hiPSC as previously described (Burridge et al., 2014). At 10 d of the differentiation, beating cardiomyocytes were switched to RPMI 1640 medium (Thermo Fisher) supplemented with B27 Supplement (Thermo Fisher). To purify the iPSC-CM population and eliminate potential contaminating mesodermal and endodermal populations, iPSC-CMs were cultured in starvation medium: RPMI 1640 without glucose (Thermo Fisher) supplemented with B27 and 213 μg/mL ascorbic acid (Sigma-Aldrich). iPSC-CMs were further expanded one passage in RPMI 1640 medium supplemented with B27 Supplement and the addition of 2 mM CHIR99021 (Tocris) per a recently described method for the proliferation of iPSC-CMs (*59*). Then cardiomyocytes were cultured in maturation medium, high in fatty acids to favor fatty acid oxidation (*60*). The maturation medium was changed every other day until experiments.

### Plasmids

All constructs were confirmed by DNA sequencing. pGEX 6P-4 vector (GE Healthcare Life Science) or pDuet1 His-HRV3C and pDuet1 His-HRV3C-AviTag were used to express GST- or His-tagged proteins, respectively, in bacteria. pEF1a-GST-6P-4 or pEF1a-GST-P-N4 vector was used to express amino terminal or carboxyl terminal GST-tagged proteins in 293T cells. pFLAG-C4 was derived from pMYC-C4 (*20*) by replacing the MYC tag with FLAG tag. This plasmid was used to transfect 293T for the overexpression of protein. pMSCV-neo MYC-C4 retroviral vector was derived from pMSCV-puro myc vector by replacing the puromycin selectable marker with a neomycin selectable marker from pEGFP-C4 plasmid (Clontech). This plasmid was used to make retroviral particles in 293T cells and to express amino terminal MYC tagged protein in mammalian cells. pInducer200 (puro) tet-on EGFP-C4 plasmid was derived from pInducer20 plasmid (Addgene #44012) by: 1) by replacing the tetracycline inducible minimal CMV promoter with the TRE3G promoter from LT3GPIR plasmid (Addgene #111177), 2) inserting EGFP cDNA from pEGFP-C4 (Clontech) plasmid in the downstream of TRE3G promoter, 3) replacing the UbC promoter with the human PGK promoter from pCW-Cas9-Blast plasmid (Addgene #83481), and 4) replacing the neomycin selection marker with the puromycin selection maker cDNA from pMSCV-puro plasmid (Clontech). This plasmid was used to make lentiviral particles in 293T cells and to express doxycycline inducible C-terminal EGFP-tagged protein in mammalian cells. pInducer200 (puro) tet-on EGFP-C4 shRNA was derived from pInducer200 (puro) tet-on EGFP-C4 by adding the miR-E sequence from LT3GPIR plasmid downstream of EGFP. This plasmid was used to make lentiviral particles in 293T cells and to express doxycycline inducible C-terminal EGFP-tagged protein with shRNA in mammalian cells. pInducer250 tet-on EGFP-N4 was derived from pInducer200 (puro) tet-on EGFP-C4 by deleting the IRES-puro sequence and moving the multiple cloning site at the 3’ end of EGFP to the 5’ end. pInducer210 (blast) tet-on mScarlet-I-N4 lentiviral vector was derived from pInducer20 plasmid by: 1) replacing the tetracycline inducible minimal CMV promoter with the TRE3G promoter, 2) inserting mScarlet-I cDNA from pmScarlet-i_C1 plasmid (Addgene #85044) downstream of the TRE3G promoter, 3) replacing the UbC promoter with the human PGK promoter, and 4) replacing the neomycin selection marker cDNA with a blasticidin selection maker cDNA from pCW-Cas9-Blast plasmid. pLKO2-hygro H1 lentiviral vector was derived from TRC2-pLKO-puro (Millipore-Sigma, SHC201) plasmid by replacing the puromycin selectable marker gene and human U6 promotor with the hygromycin selectable marker gene from pMSCV-hygro (Clontech) and the human H1 promoter from pSUPER.retro.puro (Oligoengine), respectively. This construct was used to make lentiviral particles in 293T cells and to express shRNA in mammalian cells. pLKO2-H2B-emiRFP-T2A-hygro H1 lentiviral vector was derived from pLKO2-hygro H1 plasmid by inserting the H2B-emiRFP sequence from pH2B-emiRFP670 (Addgene, #136571) with a T2A sequence at the 5’ end of the hygromycin selectable marker gene. pEGFP-C4, pmScarlet-i_C1 (Addgene, #85044), and piRFP670-N1 (Addgene, #45457) were used to express EGFP, mScarlet-I, and iRFP670 tagged proteins, respectively, in mammalian cells.

Human nesprin-2 (SYNE2-5) SR11 (1423-1531 aa), SR12 (1532-1649 aa), SR13 (1650-1745 aa), SR11-12 (1423-1649 aa), SR12-13 (1532-1745 aa), and SR11-13 (1423-1745 aa), SR51-54 (6021-6574 aa), and human zyxin-LIM (329–572) cDNAs were obtained by PCR from HeLa cell mRNA. Human nesprin-2 D1625A and D1629A mutations were previously described (*34*). Nesprin-2 cDNA was inserted into a vector with NotI restriction sites. zyxin-LIM cDNA was inserted into a vector with BamHI and NotI restriction sites. cDNAs for human FHOD1 and its variants, mouse Dia1, and human profilin1 and 2a were previously described (*20*). Human FHOD3-1 cDNA was obtained from transOMIC Technology Inc. cDNAs for human FHOD3-3 and human alpha-actinin-2 were obtained by PCR from deidentified human heart tissue (Columbia Heart Bank). All candidate HCM-associated FHOD3-3 mutations were previously described and introduced in FHOD3-3 cDNA. FHOD3-3 R124A (AGA to GCA) R125A (AGG to GCG), S397D (AGC to GAC) S521D (AGT to GAC), I1163A (ATT to GCT), S1589D (TCC to ATC) S1595D (TCT to GTC) T1599D (ACC to ATC), S1589A (TCC to AGC) S1595A (TCT to GGC) T1599A (ACC to AGC), and shRNA resistance (3796bp ATAGACCAGTTGGAGAACAAT to AT***C***GA***T***CAG***C***TGGA***A***AA***T***AA***C***) mutations were generated by PCR. All FHOD3 cDNAs were inserted in a vector by NotI restriction site.

The shRNA sequences were: for NC (Sigma, 5’-CAACAAGATGAAGAGCACCAA-3’), NTC (5’-ATTAAATAACTACTGACGTCCG-3’), and FHOD1 (Sigma, 5’-CTGGACATGACAGCGACATGC-3’). The shRNA sequences for mouse nesprin-2 and human FHOD3 were previously described (*37, 61*).

### Primary antibodies

Rabbit-raised polyclonal GFP (IF: 1:400; ThermoFisher, A-11122); mouse-raised monoclonal GFP (WB: 1:1000; Santa Cruz, sc-9996); mouse-raised monoclonal FLAG (WB: 1:2000; Millipore Sigma, F1804-50UG); rabbit-raised polyclonal MYC (WB: 1:4000 and IF: 1:400; Cell Signaling, 2278S); mouse-raised monoclonal pericentrin (IF: 1:400; BD Biosciences, 611814); mouse-raised monoclonal β-catenin (IF: 1:400; ThermoFisher, 13-8400); rat-raised monoclonal α-tubulin (WB: 1:2000 and IF: 1:40; European Collection of Authenticated Cell Cultures, 92092402); mouse-raised monoclonal GAPDH (WB: 1:5000; Abcam, ab8245); mouse-raised monoclonal cardiac Troponin T (IF: 1:400; BD Biosciences, 564766); rabbit-raised polyclonal zyxin (WB: 1:2000; Cell Signaling, 3553S); rabbit-raised polyclonal nesprin-2 (WB:1:200, a gift from Dr. Didier Hodzic at Washington University in St. Louis); rabbit-raised polyclonal FHOD1 (WB: 1:1000; Santa Cruz, sc-99209, discontinued); and rabbit-raised polyclonal FHOD3 (WB: 1:2000; Millipore Sigma, HPA024696-100UL). Note IF is for immnunofluorescent staining and WB is for Western blotting.

### Virus production and infection

293T cells were transfected with retro or lentiviral vectors and VSV-G pseudotype packaging plasmids. Medium containing the produced virus was harvested at 24, 30, 36, 48 h after transfection. After clarifying cell debris, the medium containing the virus was stored at -80 °C. To change the medium and concentrate the virus solution, virus particles were pelleted using a SW28 rotor (Beckman Coulter) at 25,000 rpm for 90 min at 4 °C. The virus was resuspended in DMEM, and the 100X concentrated virus solution was frozen at -80 °C. For NIH3T3 infection, virus solution was added to cells in the presence of 2 μg/ml polybrene (Millipore) and incubated for 1 d. For infecting iPSC-CM, the concentrated virus solution was added to cells in the presence of 2 μg/ml polybrene and incubated for 2 d.

### Protein production and purification

For production of proteins in mammalian cells, 293T cells on 150 mm plates were transfected with 30 μg of plasmid DNA with calcium phosphate precipitation method for 6 h. Two days after transfection, the cells were lysed with mammalian GST lysis buffer (25 mM Tris pH 7.4, 150 mM NaCl, 5 mM EDTA, 1% Triton-X, 10% glycerol, 1 mM DTT, 1 mM phenylmethylsulfonyl fluoride (PMSF)), and the soluble fraction was used for the protein purification. All other GST-tagged proteins were expressed in BL21(DE3) *E.coli* strain, and the bacteria after the induction of protein expression was lysed by sonication with bacteria GST lysis buffer (50 mM Tris-HCl pH=8.0, 150 mM NaCl, 0.05% Triton-X, 10% glycerol, 2 mM β-mercaptoethanol). The soluble fraction was used for the protein purification. GST-proteins were immobilized on GSH-Sepharose beads (Cytiva). His-tagged proteins were purified after expression in LOBSTR *E.coli* strain (Kerafast). The bacteria were lysed by sonication with bacteria His lysis buffer (50 mM Na_2_HPO_4_ pH=8.0, 400 mM NaCl, 40 mM immidazole, 0.02% Triton-X, 10% glycerol, 2 mM β-mercaptoethanol). The soluble fraction was used for the protein purification. His-tagged proteins were immobilized on Ni-NTA agarose beads (Qiagen). Immobilized proteins were cleaved with Turbo3C protease. The released and cleaved proteins were run on a PD-10 column (Cytiva) in phosphate-buffered saline (PBS) containing 10% glycerol, 0.02% Triton-X, and 1 mM DTT and concentrated by Amicon concentrator (Millipore). The proteins were stored at -80 °C. Purified FHOD3 protein was dephosphorylated with ʎ protein phosphatase (NEB) according to the manufacture’s protocol.

### Analytical Protein Assays

BLI interaction assay was previously described (*34*). Thermal stability assay was done using a Prometheus nanoDSF instrument (NanoTemper Technologies). Data from these results were plotted using Prism software (GraphPad Software).

### Actin assays

For the F-actin binding assay (high speed pelleting assay), 16 µM rabbit skeletal muscle actin was incubated in G-buffer (5 mM Tris-HCl pH 8.0, 0.2 mM CaCl_2_, 0.2 mM ATP, and 0.5 mM DTT) at 4 °C for at least 1 h. The G-actin was spun down at 100,000 x *g* for 20 min. The supernatant containing the G-actin was adjusted to F-buffer (50 mM KCl, 2 mM MgCl_2_, and 1 mM ATP pH 7.4) and incubated at 24°C for 2 h at 8 μM concentration. Proteins to be tested were preincubated in F-buffer at 4°C for 30 min.

Insoluble material was removed by centrifugation at 100,000 x *g* for 20 min at 4°C and the proteins mixed with the F-actin solution at 1:1 ratio resulting in 4 μM F-actin. The solution was incubated at 24°C for 60 min and spun down at 100,000 x *g* for 20 min at 4°C. The supernatant was mixed with SDS sample buffer. The pellet was washed with F-buffer once and resuspended in SDS sample buffer. Proteins in these fractions were separated by SDS-PAGE and stained with SimplyBlue SafeStain.

For the actin bundling assay (low speed pelleting assay), G-actin was polymerized in KMEI buffer (50 mM KCl, 1 mM MgCl_2_, 1 mM EGTA, and 10 mM imidazole pH 7.0) at 24 °C for 2 h. After the incubation, unlabeled phalloidin (actin:phalloidin = 1:2) was added and incubated for 5 min. Proteins to be tested were preincubated in KMEI buffer at 4 °C for 30 min. The F-actin and the protein solutions were clarified by centrifugation at 16,000 x *g* at 4 °C for 5 min and the supernatants mixed at a 1:1 ratio resulting 2 μM F-actin. The solution was incubated at 24°C for 15 min and spun down at 16,000 x *g* for 5 min at 4°C. The supernatant was mixed with SDS sample buffer. The pellet was washed with KMEI buffer once and resuspended in SDS sample buffer. Proteins in the samples were separated by SDS-PAGE and stained with SimplyBlue SafeStain.

For the fluorescent actin bundling assay, polymerized actin was incubated with Alexa Fluor 488 conjugated phalloidin (actin:phalloidin:488-phalloidin = 2:1:1) for 5 min and then incubated with proteins (as above) to test for bundling. After incubation, the bundled F-actin solution was diluted in 20 times with KMEI buffer and applied to 0.01 % poly-L-lysine coated coverslips. The samples were imaged by TIRF microscopy with the microscope described.

For the pyrene actin polymerization assay, proteins were incubated in 2X KMEI buffer at 37 °C for 15 min. Pyrene-labeled depolymerized actin (4 μM rabbit skeletal muscle actin or 8 μM human non-muscle actin, 10% pyrene labeled) in G-buffer was incubated with the indicated profilin at 37 °C for 5 min. The actin and test protein solution were mixed resulting in 2 μM muscle or 4 μM non-muscle actin. Pyrene actin fluorescent intensity was measured by SpectraMax i3x (Molecular Devices) every 10 s for 45 min.

### Western blotting and GST-pulldown interaction assay

For western blotting, proteins suspended in SDS sample buffer were separated by SDS-PAGE. The proteins were transferred to nitrocellulose, probed with indicated antibodies and then detected either by infra-red fluorescence with Odyssy CLx (LI-COR Inc.). For GST-pulldown, lysates in KLB were prepared from transfected 293T cells, clarified by centrifugation and then incubated with GST-protein immobilized beads at 4 °C for 1.5 hr. The beads were then washed four times with kinase lysis buffer (KLB: 25 mM Tris-HCl 7.4, 150 mM NaCl, 5 mM EDTA, 1% Triton X-100, 10 mM sodium pyrophosphate, 10 mM β-glycerophosphate, 10 mM NaF, 1 mM sodium orthovanadate, 10% glycerol, and 1 mM phenylmethylsulfonyl fluoride) and bound proteins eluted by boiling in SDS sample buffer.

### LPA stimulation

A day before serum-starvation, NIH3T3 fibroblasts were plated either on acid-washed coverslips for indirect immunofluorescence staining or on 20 mm diameter bottom coverslip dishes (Mattek). The next day, cells at about 40% confluency on coverslips were washed three times with PBS, three times with DMEM, and then DMEM containing 10 mM HEPES pH 7.4 and 0.1% (v/v) fatty acid free bovine serum albumin was added. For indirect immunofluorescence staining, cells were serum-starved for 2 d. Then, the cell monolayer was wounded with a P200 pipet tip. After 1 h, the cells were stimulated with 10 μM LPA and fixed at the indicated times. In experiments expressing FHOD3 proteins in NIH3T3 fibroblasts, cells were pretreated with 1 μM AZD for 30 min before LPA stimulation because FHOD3 was hyperphosphorylated upon expression in NIH3T3 fibroblasts. This reduced FHOD3 activity required for nuclear movement (see ref. 20). For live imaging, cells were serum-starved for 2 d. The night before stimulating the cells with 20 μM LPA, 100 ng/ml doxycycline hyclate (Sigma-Aldrich) was added. One hour after wounding, the cells were mounted on a microscope and stimulated with LPA.

### Fluorescence Microscopy

For indirect immunofluorescence microscopy, cells were fixed with 4% paraformaldehyde in PBS for 20 min, permeabilized with 0.1% Triton-X, and then blocked with PBS containing 0.1% Triton-X and 1% BSA for 30 min. The cells were labeled with primary antibodies and then fluorescently labeled secondary antibodies, phalloidin, and DAPI. TIRF images were acquired with a 60× PlanApo TIRF objective (NA 1.49) and an ORCA ERI CCD camera (Hamamatsu) equipped with TIRF illuminator and fiber-optic-coupled lasers on a Nikon Eclipse Ti microscope controlled by Nikon’s NIS-Elements software. Confocal images were acquired with a 60× PlanApo TIRF objective (NA 1.49) and ORCA-Fusion BT CMOS CCD camera (Hamamatsu) equipped with X-Light V3 Spinning Disk (CrestOptics) mounted on inverted Ti2 microscope (Nikon) controlled by NIS-Elements software.

### Transfection and replating of cardiomyocytes

iCell® Cardiomyocytes2 were transfected with ViaFect reagent (Qiagen) according to the manufacture’s protocol. Two days after the transfection, the cells were dissociated and replated on gelatin-coated 7 mm diameter coverslip dishes (Mattek) according to the FUJI Cellular Dynamics protocol. For replating iPSC-CM, cells were incubated with TrypLE select (ThermoFisher Scientific) at 37 °C until they were fully detached from the plate. The cells were resuspended by adding an equal volume of KnockOut™ Serum Replacement (KOSR) medium (ThermoFisher Scientific) and then spun down at 200 x *g* for 5 min. Cell pellets were resuspended in a medium of 90% of Maturation Medium with high fatty acids, 10% KOSR medium, and 5 μM Y-27632 dihydrochloride and plated on Matrigel-coated, 96-well glass bottom plates. The next day, the medium was replaced with Maturation Medium high in fatty acids. In 3 d, the cells were fixed. For doxycycline inducible FHOD3 expression, cells were cultured in the presence of 100 ng/ml doxycycline hyclate (Sigma-Aldrich) continuously until the cells were fixed or lysed.

### Generation and maintenance of Fhod3 R611P mouse

The Institutional Animal Care and Use Committee at Columbia University Irving Medical Center approved all protocols. A point mutation (AGG to CCC; R611P) was introduced into exon 14 of the *Fhod3* gene, and an FNF (Frt-Neo-Frt) cassette was inserted into the intron downstream of exon 14 to generate the *Fhod3* R611P mouse line. A 748 bp left homology arm (LHA) containing exon 14 with the R611P mutation and a 622 bp right homology arm (RHA) were prepared via DNA synthesis and cloned into the pUC57-FNF vector to construct the pUC57-FNF-*Fhod3*R611P gene-targeting vector. This vector was linearized and electroporated into KV1 (129B6N) embryonic stem (ES) cells. The FNF cassette conferred G418 resistance during the gene-targeting process in these cells.

Targeted ES cell clones were identified and injected into C57BL/6J blastocysts to generate chimeric mice. Male chimeras were bred with B6.Cg-Tg(ACTFLPe)9205Dym/J mice (The Jackson Laboratory; stock no. 005703) to remove the Neo cassette, producing heterozygous *Fhod3* (R611P/+) mice with a remaining Frt site downstream of exon 14. Offspring were backcrossed to wild-type C57BL/6J mice for several generations to establish a >90% pure C57BL/6J background. Finally, heterozygous males and females were intercrossed to produce *Fhod3*^R611P/R611P^, *Fhod3*^R611P/+^, and *Fhod3*^+/+^ mice used in experiments.

Mice were housed in a barrier facility under a 12-hour light/dark cycle, provided a chow diet, and genotyped by PCR using genomic DNA extracted from tail clippings.

### Mouse Studies

All animal procedures were approved by the Institutional Animal Care and Use Committee of Columbia University Medical Center and conformed to the NIH *Guide for the Care and Use of Laboratory Animals*. Fhod3 heterozygous mutant mice (C57BL/6 background; 25–30 g; age 15–30 weeks) and wild-type littermates were used. Mice were randomized into four groups (n = 5–8/group): WT + Vehicle, WT + Angiotensin II (Ang II), Fjhod3 Mutant + Vehicle, and Fhod3 Mutant + Ang II. Ang II (2.5 mg/kg/day) or vehicle (0.9% saline) was administered via subcutaneously implanted Alzet osmotic mini pumps (Model 1004; Durect Corp.) for 28 d. Pumps were placed in subcutaneous pockets between the scapulae.

At day 28, mice were anesthetized with isoflurane (2%) and subjected to transthoracic echocardiography (Vevo 3100, VisualSonics; 30-MHz probe).

### Tissue Collection and Histological Analysis

One day following imaging, mice were weighed, euthanized, and hearts excised. Hearts were either flash-frozen in liquid nitrogen for molecular analyses or fixed in 4% paraformaldehyde. For histology, hearts were paraffin-embedded, sectioned at 4 μm, and stained with Masson’s Trichrome. Sections were evaluated under 400× magnification (Leica Microsystems).

### RNA Isolation and qRT-PCR

Total RNA was extracted from heart tissue using TRIzol reagent (Invitrogen) and treated with DNase. cDNA synthesis was performed using the ProtoScript II First-Strand cDNA Synthesis Kit (New England Biolabs). Quantitative real-time PCR was conducted using SYBR Select Master Mix (Applied Biosystems) on the ABI 7300 Fast System. Transcript levels were normalized to 18S rRNA using the 2^−ΔΔCt method.

### Quantification and Statistical Analysis

For F-actin bundling and binding quantification, stained protein bands on the gel were scanned and quantified using ImageJ software and plotted on the graph using Microsoft Excel software. For F-actin strain quantification, the number of actin cables that exhibited elevated EGFP-zyxin-LIM signals at thinned actin bundle sites was quantified from fixed images. For the centrosome reorientation assays, the position of centrosome relative to the axis between the nuclei and the leading edge was analyzed from images of DAPI and tubulin and/or β-catenin/pericentrin antibody-labeled cells as previously described (*44, 62*). Nuclear and centrosomal positions relative to the cell centroid in NIH3T3 fibroblasts were determined from images using Cell Plot software (*21*). Heart weight (HW) and tibia length (TL) were measured to calculate HW/TL and HW/body weight (BW) ratios. Left ventricular mass (LVM) was measured by echocardiography using the Devereux formula. Electrocardiographic recordings were obtained simultaneously using a four-limb lead setup and B08 amplifier (Emka Technologies). Data were analyzed using IOX Software v1.8.9.18 and ECG Auto v1.5.12.50, with interval measurements averaged from three consecutive cardiac cycles. Fibrosis was quantified using ImageJ software (NIH, https://imagej.nih.gov/ij/).

Statistical evaluation of strained F-actin in NIH3T3 fibroblasts (Fig. 3B, D, and F), the position of the nucleus and centrosome in NIH3T3 fibroblasts (Fig. 5B and fig. S5E), and mouse analysis (Fig. 6 and fig. S8) was assessed by one-way ANOVA followed by Tukey’s multiple comparison test using Prism software (GraphPad Software). Statistical evaluation of centrosome reorientation (Fig. 5B and fig. S5E) and Mendelian ratio of *Fhod3* genotype (fig. S8A) was assessed by Chi-square test using Prism software.

## Supporting information

Suppelemental Figures

## ACKNOWLEDGEMENTS

Research reported in this publication was supported by NIH grants R35 GM136403 to GGG and RO1 HL159389 to HJW and GGG. The *Fhod3*^R611P/+^ mouse was made with generous support of Columbia Precision Medicine Pilot Grant Program and utilized the facilities of Genetically Modified Mouse Models Shared Resource core at Columbia University Herbert Irving Comprehensive Cancer Center. The content is solely the responsibility of the authors and does not necessarily represent the official views of the NIH.

## FIGURE LEGENDS

**Supplemental Figure 1.** The biophysical properties of the interaction between SR11-13 of nesprin-2G and SRBD-DID region of FHOD1. **A**. Complex structure of FHOD1 SRBD-DID with nesprin-2G SR11-12 (PDB:6XF1) superposed with predicted AlphaFold3 model of FHOD1 SRBD-DID with the extended nesprin2G SR11-13 fragment. SR11-12 shown in gray, SR11-13 in blue and FHOD1 SRBD-DID in orange. Interaction between FHOD1 SRBD-DID and nesprin-2G is unchanged when the two structures are compared. **B**. Thermal stability assay of nesprin-2G SR11-12 vs. SR11-13. Top panels show the actual measurement, lower panels the first derivative, to identify the melting temperature Tm. Nesprin-2 SR11-13 shows a Tm of 60 °C compared to 46 °C measured for nesprin-2G SR11-12, indicating increased stability. Dotted line represents the Tm for the proteins. C. BLI analysis of FHOD1 SRBD-DID with nesprin-2G SR11-12 and nesprin-2G SR11-13. The nesprin-2G fragments were immobilized as ligand and tested against FHOD1 SRBD-DID at 5 concentrations (40, 80, 160, 320, 640 nM). Red lines represent the fitted curves.

**Supplemental Figure 2.** The interaction between SR11-13 of nesprin-2G and SRBD region of FHOD1/3. **A**. 293T cells were transfected with a plasmid driving the expression of FLAG-tagged FHOD1/3. The transfected cells were lysed and subjected for GST-pulldown assay with the indicated nesprin-2G constructs. The GST-pulldown products and cell lysates were subjected to SDS-PAGE and immunoblotting. The images indicate Ponceau S staining and immunoblotting with the indicated antibody. The number on the left side of gel indicated molecular weight in kDa. **B**. TIRF image of fluorescently labeled F-actin from F-actin bundling assay. F-actin decorated with fluorescent phalloidin was incubated with indicated proteins. The solution was put onto poly L-lysine coated coverslips and image was taken with TIRF microscope. Bar, 5 μm. C. The image of Coomassie Brilliant Blue stained acrylamide gel from *In vitro* F-actin bundling assay. The indicated purified proteins were incubated with F-actin and spun down at 16,000 xg. The supernatant and pellet fractions were subjected for SDS-PAGE and the gel was stained by Coomassie Brilliant Blue solution. Intended input amount of protein is indicated. The number on the left side of gel indicated molecular weight in kDa.

**Supplemental Figure 3.** FHOD1 F-actin bundling activity is specifically activated by Nesp2 SR11-13. **A, B, and C**. The image of Coomassie Brilliant Blue stained acrylamide gel from *In vitro* F-actin bundling assay. The indicated purified proteins were incubated with F-actin and spun down at 16,000 xg. The supernatant and pellet fractions were subjected for SDS-PAGE and the gel was stained by Coomassie Brilliant Blue solution. Intended input amount of protein is indicated. The number on the left side of gel indicated molecular weight in kDa.

**Supplemental Figure 4.** Nesprin-2 SR11-13 is the unique activator of FHOD1/3 and specifically activates their F-actin bundling activity. The image of Coomassie Brilliant Blue stained acrylamide gel from *In vitro* F-actin bundling assay. The indicated purified proteins were incubated with F-actin and spun down at 16,000 xg. The supernatant and pellet fractions were subjected for SDS-PAGE and the gel was stained by Coomassie Brilliant Blue solution. Intended input amount of protein is indicated. The number on the left side of gel indicated molecular weight in kDa. **B**. The graph of F-actin bundling efficiency by actin binding protein (ABP). The amount of actin and ABP was quantified from the gel in **A**. These were plotted on a graph. **C**. *In vitro* F-actin polymerization Assay using muscle actin. 2 μM muscle actin (10% pyrene-labeled muscle actin) was pre-incubated with 4 μM of profilin1 at 37 °C in a buffer maintaining actin in monomer state. Then, the profilin1-actin was mixed with indicated protein in an actin polymerization buffer at 37 °C, and the fluorescent absorbance for pyrene-actin is measured over time. These were plotted on a graph. The concentration of protein in the reaction is 1 μM Nesp2 SR11-13, 200 nM of FHOD1, and 20 nM mDia1. **D**. *In vitro* F-actin polymerization Assay using non-muscle actin. 4 μM non-muscle actin (10% pyrene-labeled muscle actin) was pre-incubated with 8 μM of profilin1 at 37 °C in a buffer maintaining actin in monomer state. Then, the profilin1-actin was mixed with indicated protein in an actin polymerization buffer at 37 °C, and the fluorescent absorbance for pyrene-actin is measured over time. These were plotted on a graph. The concentration of protein in the reaction is 2 μM Nesp2 SR11-13, 400 nM of FHOD1, and 40 nM mDia1.

**Supplemental Figure 5.** FHOD1 knockdown cells upregulates strain of F-actin over nucleus and overexpression of zyxin rescues nuclear movement defect of FHOD1 knockdown cells. **A-C**. Western blots of NIH3T3 fibroblasts **A**) stably expressing doxycycline inducible EGFP-zyxin-LIM and indicated shRNA , **B**) stably expressing scrambled shRNA (NC) or shRNA targeting FHOD1, doxycycline inducible EGFP-zyxin-LIM, and the indicated FHOD1 rescue constructs, and **C**) stably expressing scrambled shRNA (shNC) or shRNA targeting FHOD1 (shFHOD1) with EGFP or EGFP-zyxin. Antibodies used to probe the blots are indicated at the right. Migration of molecular mass standards (kDa) is indicated at the left. **D**. Images of LPA-stimulated wound-edge NIH3T3 fibroblasts from **C** stained for the indicated proteins and DAPI. Dashed lines show the middle line of cell. Bar, 10 μm. **E**. Histogram of centrosome and nuclear positions relative to the cell centroid (defined as “0”; negative values toward the cell rear). Centrosome orientation (heat map below the histogram, values mean % of cells) toward the front of the cell for cells treated as in **D**. Values are means ± SEM; n, cells examined. *^1^ and *^3^ indicates p < 0.05 compared to nuclear position and centrosome orientation, respectively, of shNC NIH3T3 fibroblast expressing EGFP. *^2^ and *^4^ indicates p > 0.05 compared to nuclear position and centrosome orientation, respectively, of shFHOD1 NIH3T3 fibroblast expressing EGFP.

**Supplemental Figure 6.** FHOD3 is required for sarcomere formation and maintenance. **A**. The FHOD3 expression in iPSC-CM expressing scrambled shRNA or the one targeting FHOD3. iPSC-CM was infected with lentivirus expressing indicated shRNA and H2B-emiRFP670. The cell lysate was subjected with immunoblotting. Antibodies used to probe the blots are indicated at the right. Migration of molecular mass standards (kDa) is indicated at the left. **B**. Images of iPSC-CM expressing scrambled shRNA or the one targeting FHOD3. The cells were replated on glass-bottom dish for 4 days. Then, the cells were fixed and stained as indicated. The images were taken by a spinning disk confocal microscope. The image is the maximum projection image from Z-stack images. Bar, 10 μm.

**Supplemental Figure 7.** *Fhod3* mutant mouse model design and genotyping. **A**. *Fhod3* mutant mouse model knock-in design. **B**. A representative image of *Fhod3* genotyping PCR. **C**. Representative image of Sanger DNA sequencing of WT and *Fhod3* mutant mice.

**Supplemental Figure 8.** Baseline characterization of *Fhod3* mutant mouse model. **A.** Mendelian ratio of the genotypes of litters from *Fhod3* heterozygote vs heterozygote mating. **B**. Representative images of embryos at day 10.5. Bar, 1 mm. **C**. Representative image of Fhod3 protein expression in the ventricle (n=5). Antibodies used to probe the blots are indicated at the right. Migration of molecular mass standards (kDa) is indicated at the left. **D**. Body weigh measurements (n=10-20). **E**. Left ventricle ejection fraction. **F**. Heart weight normalized to body weight (n=4-5). **G**. Left ventricle mass normalized to body weight (n=4-5).

**Supplemental Figure 9.** Echocardiographic images of Ang II-treated *Fhod3* mutant mouse. **A**. Representative image of the LV wall thickness. **B**. Representative image of the interventricular septum and the Left atrial.

**Supplemental Movie 1**. Nuclear movement strains F-actin cables over nucleus. **A**. The time lapse images of nuclear movement with F-actin strain reporter, EGFP-zyxin-LIM. Wounded monolayer of serum-starved NIH3T3 cells stably expressing H2B-emiRFP670 and doxycycline-inducible EGFP-zyxin-LIM and Lifeact-mScarlet-I were stimulated with LPA. The images of cells were started to be taken from 15 minutes before the stimulation and to 2 hours and 15 minutes after the stimulation for every 2 minutes and 30 seconds by spinning disk confocal microscope. The dorsal (bottom) and ventral (top) portions of image were obtained from maximum Z-projection images from corresponding Z stacks. Red, green, and blue colors show F-actin (Lifeact-mScarlet-I), zyxin-LIM (EGFP-zyxin-LIM), and nucleus (H2B-emiRFP670). **B**. The magnified time-lapse images in the area over the nucleus from **A**.

**Supplemental Movie 2**. FHOD3 and sarcomere formation. **A**. The images of iPSC-CM expressing scrambled shRNA (2A-1) or the one targeting FHOD3 (2A-2). iPSC-CM was infected with lentivirus expressing indicated shRNA and H2B-mScarlet-I. After 5 days of infection, time-lapse images of cells by phase and epifluorescence (Red, H2B-mScarlet-I, nucleus) were taken at 3 frames per second for 10 seconds. **B**. The time-lapse images of sarcomere formation after replating iPSC-CM cells. iPSC-CM was transfected with plasmids expressing Lifeact-EGFP, mScarlet-I-alpha-actinin, and iRFP670-NLSx3. The cells were replated and the time-lapse images of cells were taken every 5 minutes by epifluorescence microscopy. 2B-1 shows mScarlet-I-α-actinin (green, sarcomere)/ iRFP670-NLSx3 (red, nucleus), and 2B-2 shows Lifeact-EGFP (green, F-actin)/ iRFP670-NLSx3 (red, nucleus). The time after replating is indicated at the right-bottom.

## Notes

### Competing Interest Statement

The authors have declared no competing interest.

## REFERENCES

1. J. Lammerding, Mechanics of the nucleus. Compr Physiol 1, 783–807 (2011).

2. T. P. Lele, R. B. Dickinson, G. G. Gundersen, Mechanical principles of nuclear shaping and positioning. J Cell Biol 217, 3330–3342 (2018).

3. W. Chang, H. J. Worman, G. G. Gundersen, Accessorizing and anchoring the LINC complex for multifunctionality. J Cell Biol 208, 11–22 (2015).

4. D. A. Starr, H. N. Fridolfsson, Interactions between nuclei and the cytoskeleton are mediated by SUN-KASH nuclear-envelope bridges. Annu Rev Cell Dev Biol 26, 421–444 (2010).

5. C. Guilluy et al., Isolated nuclei adapt to force and reveal a mechanotransduction pathway in the nucleus. Nat Cell Biol 16, 376–381 (2014).

6. I. L. Ivanovska, M. P. Tobin, T. Bai, L. J. Dooling, D. E. Discher, Small lipid droplets are rigid enough to indent a nucleus, dilute the lamina, and cause rupture. J Cell Biol 222, (2023).

7. S. Antoku, T. U. Schwartz, G. G. Gundersen, FHODs: Nuclear tethered formins for nuclear mechanotransduction. Front Cell Dev Biol 11, 1160219 (2023).

8. S. Kutscheidt et al., FHOD1 interaction with nesprin-2G mediates TAN line formation and nuclear movement. Nat Cell Biol 16, 708–715 (2014).

9. G. W. Luxton, E. R. Gomes, E. S. Folker, E. Vintinner, G. G. Gundersen, Linear arrays of nuclear envelope proteins harness retrograde actin flow for nuclear movement. Science 329, 956–959 (2010).

10. G. W. Luxton, E. R. Gomes, E. S. Folker, H. J. Worman, G. G. Gundersen, TAN lines: a novel nuclear envelope structure involved in nuclear positioning. Nucleus 2, 173–181 (2011).

11. A. B. Chambliss et al., The LINC-anchored actin cap connects the extracellular milieu to the nucleus for ultrafast mechanotransduction. Sci Rep 3, 1087 (2013).

12. L. M. Hoffman et al., Mechanical stress triggers nuclear remodeling and the formation of transmembrane actin nuclear lines with associated nuclear pore complexes. Mol Biol Cell 31, 1774–1787 (2020).

13. P. T. Arsenovic et al., Nesprin-2G, a Component of the Nuclear LINC Complex, Is Subject to Myosin-Dependent Tension. Biophys J 110, 34–43 (2016).

14. G. W. Luxton, D. A. Starr, KASHing up with the nucleus: novel functional roles of KASH proteins at the cytoplasmic surface of the nucleus. Curr Opin Cell Biol 28, 69–75 (2014).

15. K. E. L. Garner et al., The meiotic LINC complex component KASH5 is an activating adaptor for cytoplasmic dynein. J Cell Biol 222, (2023).

16. N. Sahabandu et al., Active microtubule-actin cross-talk mediated by a nesprin-2G-kinesin complex. Sci Adv 11, eadq4726 (2025).

17. K. Burridge, C. Guilluy, Focal adhesions, stress fibers and mechanical tension. Exp Cell Res 343, 14–20 (2016).

18. J. Z. Kechagia, J. Ivaska, P. Roca-Cusachs, Integrins as biomechanical sensors of the microenvironment. Nat Rev Mol Cell Biol 20, 457–473 (2019).

19. D. Pinheiro, Y. Bellaiche, Mechanical Force-Driven Adherens Junction Remodeling and Epithelial Dynamics. Dev Cell 47, 3–19 (2018).

20. S. Antoku et al., ERK1/2 Phosphorylation of FHOD Connects Signaling and Nuclear Positioning Alternations in Cardiac Laminopathy. Dev Cell 51, 602–616 e612 (2019).

21. A. Jayo et al., Fascin Regulates Nuclear Movement and Deformation in Migrating Cells. Dev Cell 38, 371–383 (2016).

22. F. Li Mow Chee et al., Mena regulates nesprin-2 to control actin-nuclear lamina associations, trans-nuclear membrane signalling and gene expression. Nat Commun 14, 1602 (2023).

23. G. Bonne et al., Mutations in the gene encoding lamin A/C cause autosomal dominant Emery-Dreifuss muscular dystrophy. Nat Genet 21, 285–288 (1999).

24. D. Fatkin et al., Missense mutations in the rod domain of the lamin A/C gene as causes of dilated cardiomyopathy and conduction-system disease. N Engl J Med 341, 1715–1724 (1999).

25. P. Meinke et al., Muscular dystrophy-associated SUN1 and SUN2 variants disrupt nuclear-cytoskeletal connections and myonuclear organization. PLoS Genet 10, e1004605 (2014).

26. F. D. Ravera, V.; Bocchino, P. P.; Gobello, G.; Giannino, G.; Melis, D.; Brach Del Prever, G. M.; Angelini, F.; Saglietto, A.; Giustetto, C.; Gallone, G.; Pidello, S.; Cannillo, M.; Cingolani, M. M.; Deaglio, S.; Marra, W. G.; Maria De Ferrari, G.; Raineri, C., Cardiovascular Involvement in SYNE Variants: A Case Series and Narrative Review. Cardiogenetics 15, 1–18 (2025).

27. T. Arimura et al., Dilated cardiomyopathy-associated FHOD3 variant impairs the ability to induce activation of transcription factor serum response factor. Circ J 77, 2990–2996 (2013).

28. J. P. Ochoa et al., Formin Homology 2 Domain Containing 3 (FHOD3) Is a Genetic Basis for Hypertrophic Cardiomyopathy. J Am Coll Cardiol 72, 2457–2467 (2018).

29. E. C. Wooten et al., Formin homology 2 domain containing 3 variants associated with hypertrophic cardiomyopathy. Circ Cardiovasc Genet 6, 10–18 (2013).

30. J. P. Ochoa et al., Deletions of specific exons of FHOD3 detected by next-generation sequencing are associated with hypertrophic cardiomyopathy. Clin Genet 98, 86–90 (2020).

31. G. Wu et al., Variant Spectrum of Formin Homology 2 Domain-Containing 3 Gene in Chinese Patients With Hypertrophic Cardiomyopathy. J Am Heart Assoc 10, e018236 (2021).

32. A. L. Auld, E. S. Folker, Nucleus-dependent sarcomere assembly is mediated by the LINC complex. Mol Biol Cell 27, 2351–2359 (2016).

33. A. M. Fenix et al., Muscle-specific stress fibers give rise to sarcomeres in cardiomyocytes. Elife 7, (2018).

34. S. M. Lim, V. E. Cruz, S. Antoku, G. G. Gundersen, T. U. Schwartz, Structures of FHOD1-Nesprin1/2 complexes reveal alternate binding modes for the FH3 domain of formins. Structure 29, 540–552 e545 (2021).

35. A. A. Patel, Z. A. Oztug Durer, A. P. van Loon, K. V. Bremer, M. E. Quinlan, Drosophila and human FHOD family formin proteins nucleate actin filaments. J Biol Chem 293, 532–540 (2018).

36. A. Schonichen et al., FHOD1 is a combined actin filament capping and bundling factor that selectively associates with actin arcs and stress fibers. J Cell Sci 126, 1891–1901 (2013).

37. W. Chang, S. Antoku, C. Ostlund, H. J. Worman, G. G. Gundersen, Linker of nucleoskeleton and cytoskeleton (LINC) complex-mediated actin-dependent nuclear positioning orients centrosomes in migrating myoblasts. Nucleus 6, 77–88 (2015).

38. R. Zhu, S. Antoku, G. G. Gundersen, Centrifugal Displacement of Nuclei Reveals Multiple LINC Complex Mechanisms for Homeostatic Nuclear Positioning. Curr Biol 27, 3097–3110 e3095 (2017).

39. R. Takeya, H. Sumimoto, Fhos, a mammalian formin, directly binds to F-actin via a region N-terminal to the FH1 domain and forms a homotypic complex via the FH2 domain to promote actin fiber formation. J Cell Sci 116, 4567–4575 (2003).

40. M. A. Smith et al., LIM domains target actin regulators paxillin and zyxin to sites of stress fiber strain. PLoS One 8, e69378 (2013).

41. X. Sun et al., Mechanosensing through Direct Binding of Tensed F-Actin by LIM Domains. Dev Cell 55, 468–482 e467 (2020).

42. J. D. Winkelman, C. A. Anderson, C. Suarez, D. R. Kovar, M. L. Gardel, Evolutionarily diverse LIM domain-containing proteins bind stressed actin filaments through a conserved mechanism. Proc Natl Acad Sci U S A 117, 25532–25542 (2020).

43. M. Yoshigi, L. M. Hoffman, C. C. Jensen, H. J. Yost, M. C. Beckerle, Mechanical force mobilizes zyxin from focal adhesions to actin filaments and regulates cytoskeletal reinforcement. J Cell Biol 171, 209–215 (2005).

44. E. R. Gomes, S. Jani, G. G. Gundersen, Nuclear movement regulated by Cdc42, MRCK, myosin, and actin flow establishes MTOC polarization in migrating cells. Cell 121, 451–463 (2005).

45. K. Taniguchi et al., Mammalian formin fhod3 regulates actin assembly and sarcomere organization in striated muscles. J Biol Chem 284, 29873–29881 (2009).

46. O. M. Kan et al., Mammalian formin Fhod3 plays an essential role in cardiogenesis by organizing myofibrillogenesis. Biol Open 1, 889–896 (2012).

47. A. Chopra et al., Force Generation via beta-Cardiac Myosin, Titin, and alpha-Actinin Drives Cardiac Sarcomere Assembly from Cell-Matrix Adhesions. Dev Cell 44, 87–96 e85 (2018).

48. T. Ushijima et al., The actin-organizing formin protein Fhod3 is required for postnatal development and functional maintenance of the adult heart in mice. J Biol Chem 293, 148–162 (2018).

49. S. Barko et al., Characterization of the biochemical properties and biological function of the formin homology domains of Drosophila DAAM. J Biol Chem 285, 13154–13169 (2010).

50. O. Esue, E. S. Harris, H. N. Higgs, D. Wirtz, The filamentous actin cross-linking/bundling activity of mammalian formins. J Mol Biol 384, 324–334 (2008).

51. E. S. Harris, I. Rouiller, D. Hanein, H. N. Higgs, Mechanistic differences in actin bundling activity of two mammalian formins, FRL1 and mDia2. J Biol Chem 281, 14383–14392 (2006).

52. A. Junemann et al., ForC lacks canonical formin activity but bundles actin filaments and is required for multicellular development of Dictyostelium cells. Eur J Cell Biol 92, 201–212 (2013).

53. G. Machaidze et al., Actin filament bundling and different nucleating effects of mouse Diaphanous-related formin FH2 domains on actin/ADF and actin/cofilin complexes. J Mol Biol 403, 529–545 (2010).

54. A. Michelot et al., A novel mechanism for the formation of actin-filament bundles by a nonprocessive formin. Curr Biol 16, 1924–1930 (2006).

55. E. W. Miller, S. D. Blystone, The carboxy-terminus of the formin FMNL1ɣ bundles actin to potentiate adenocarcinoma migration. J Cell Biochem 120, 14383–14404 (2019).

56. J. B. Moseley, B. L. Goode, Differential activities and regulation of Saccharomyces cerevisiae formin proteins Bni1 and Bnr1 by Bud6. J Biol Chem 280, 28023–28033 (2005).

57. C. L. Vizcarra, B. Bor, M. E. Quinlan, The role of formin tails in actin nucleation, processive elongation, and filament bundling. J Biol Chem 289, 30602–30613 (2014).

58. X. H. Xue et al., AtFH8 is involved in root development under effect of low-dose latrunculin B in dividing cells. Mol Plant 4, 264–278 (2011).

59. J. W. Buikema et al., Wnt Activation and Reduced Cell-Cell Contact Synergistically Induce Massive Expansion of Functional Human iPSC-Derived Cardiomyocytes. Cell Stem Cell 27, 50–63 e55 (2020).

60. D. A. M. Feyen et al., Metabolic Maturation Media Improve Physiological Function of Human iPSC-Derived Cardiomyocytes. Cell Rep 32, 107925 (2020).

61. T. Iskratsch et al., Formin follows function: a muscle-specific isoform of FHOD3 is regulated by CK2 phosphorylation and promotes myofibril maintenance. J Cell Biol 191, 1159–1172 (2010).

62. A. F. Palazzo et al., Cdc42, dynein, and dynactin regulate MTOC reorientation independent of Rho-regulated microtubule stabilization. Curr Biol 11, 1536–1541 (2001).

