## Supplementary material for "The nucleus activates mechano-responsiveness via FHOD-associated LINC complexes": Suppelemental Figures

### Supplemental Figure 1.

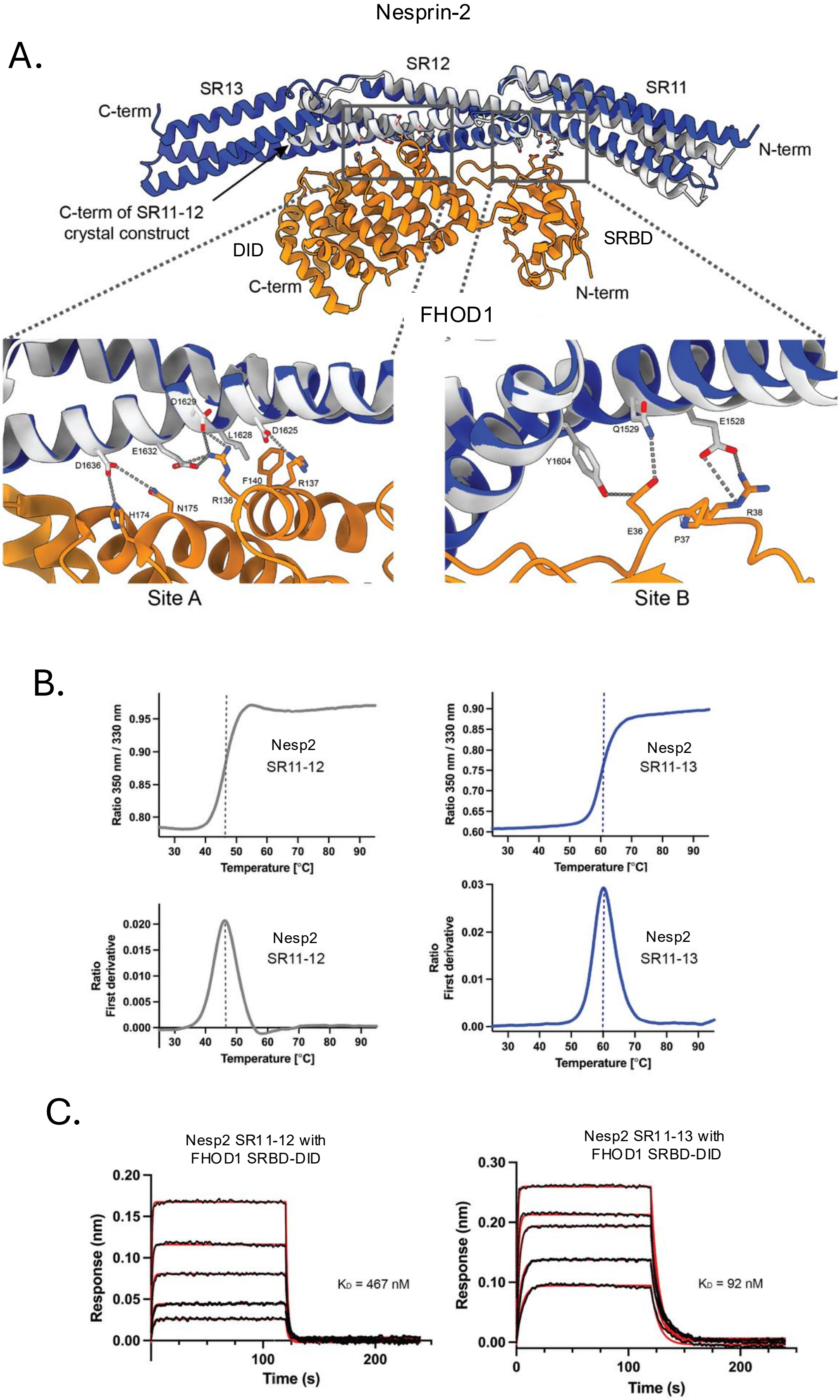

### Supplemental Figure 2.

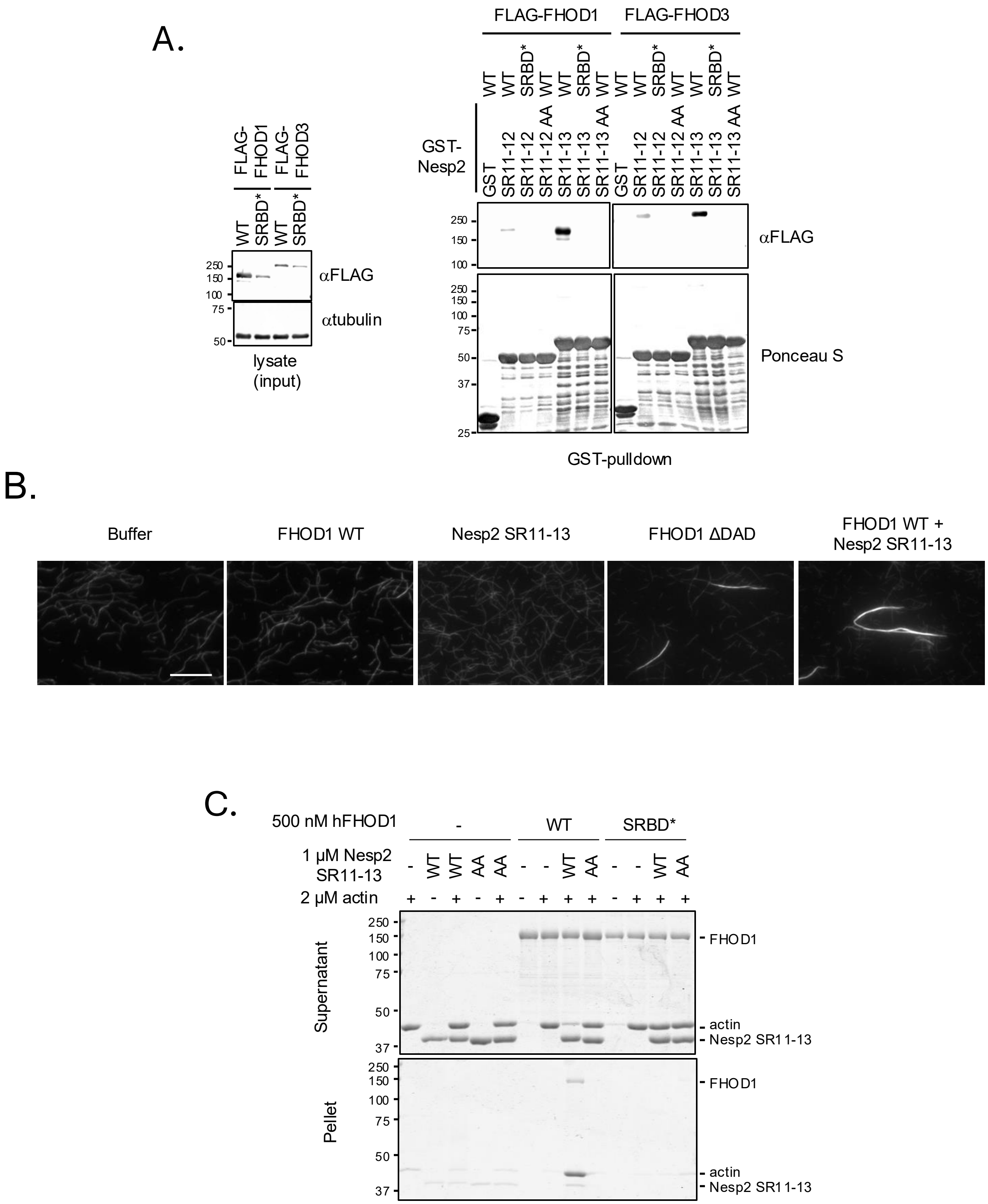

### Supplemental Figure 3.

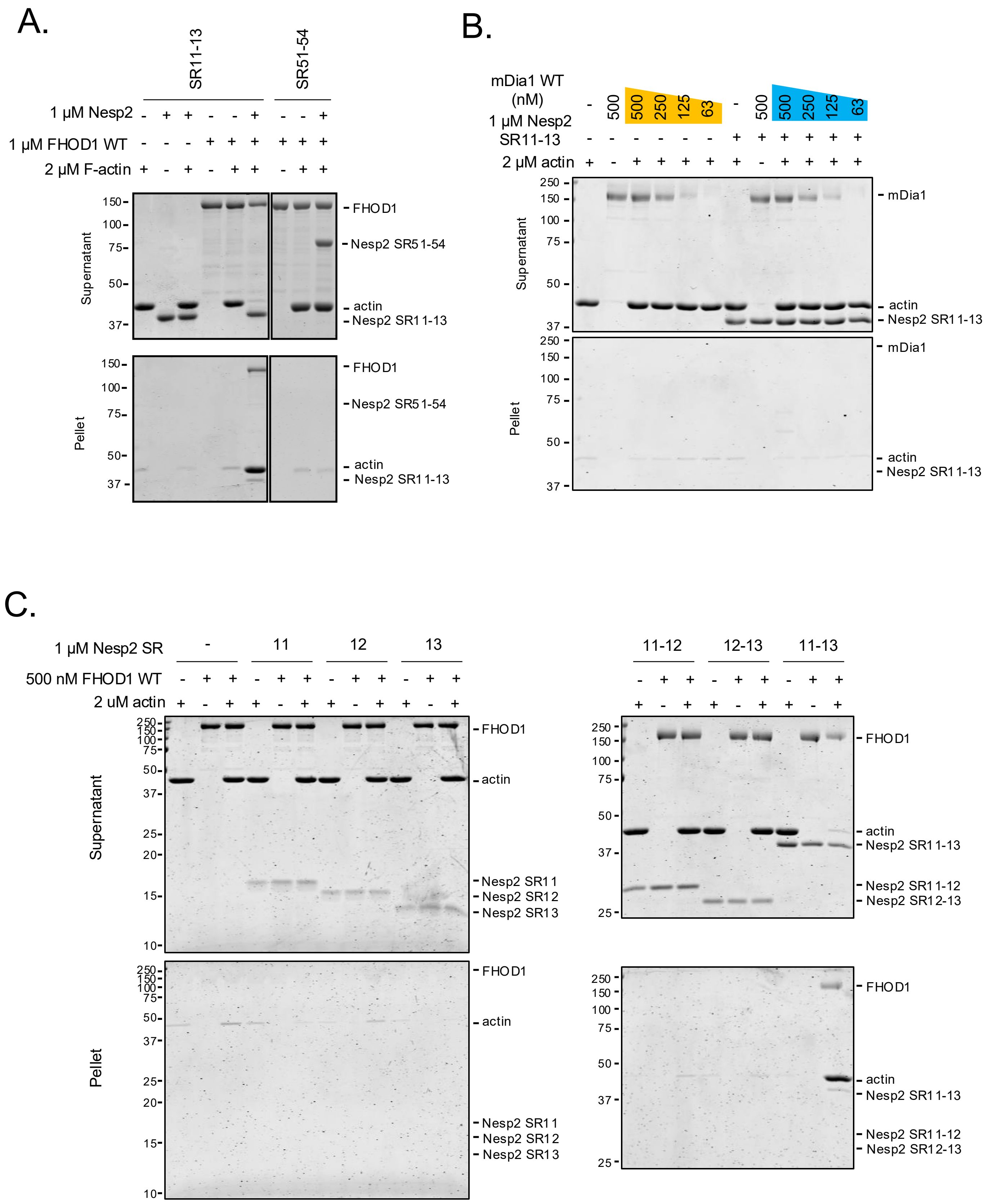

Supplemental Figure 4.

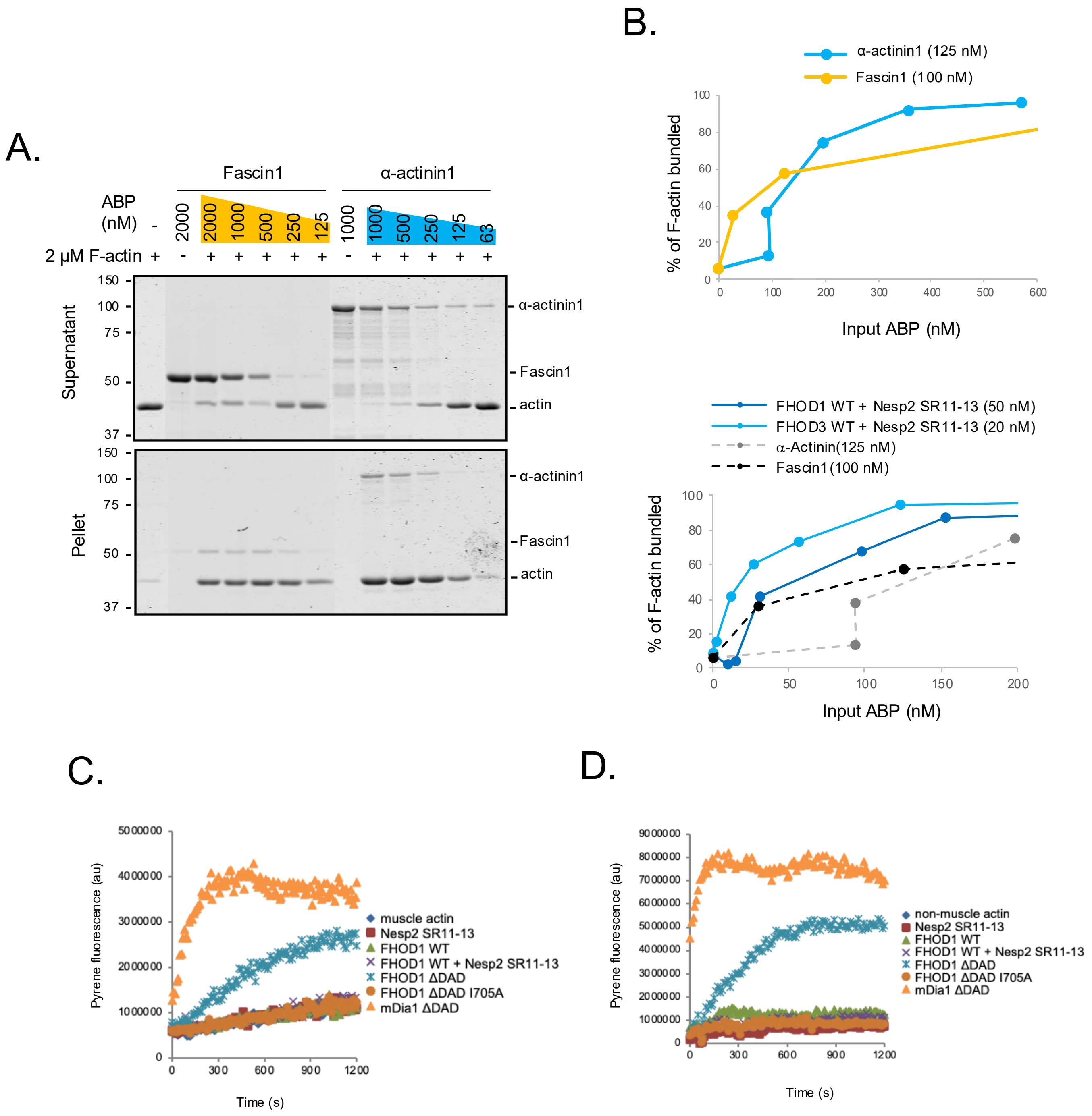

Supplemental Figure 5.

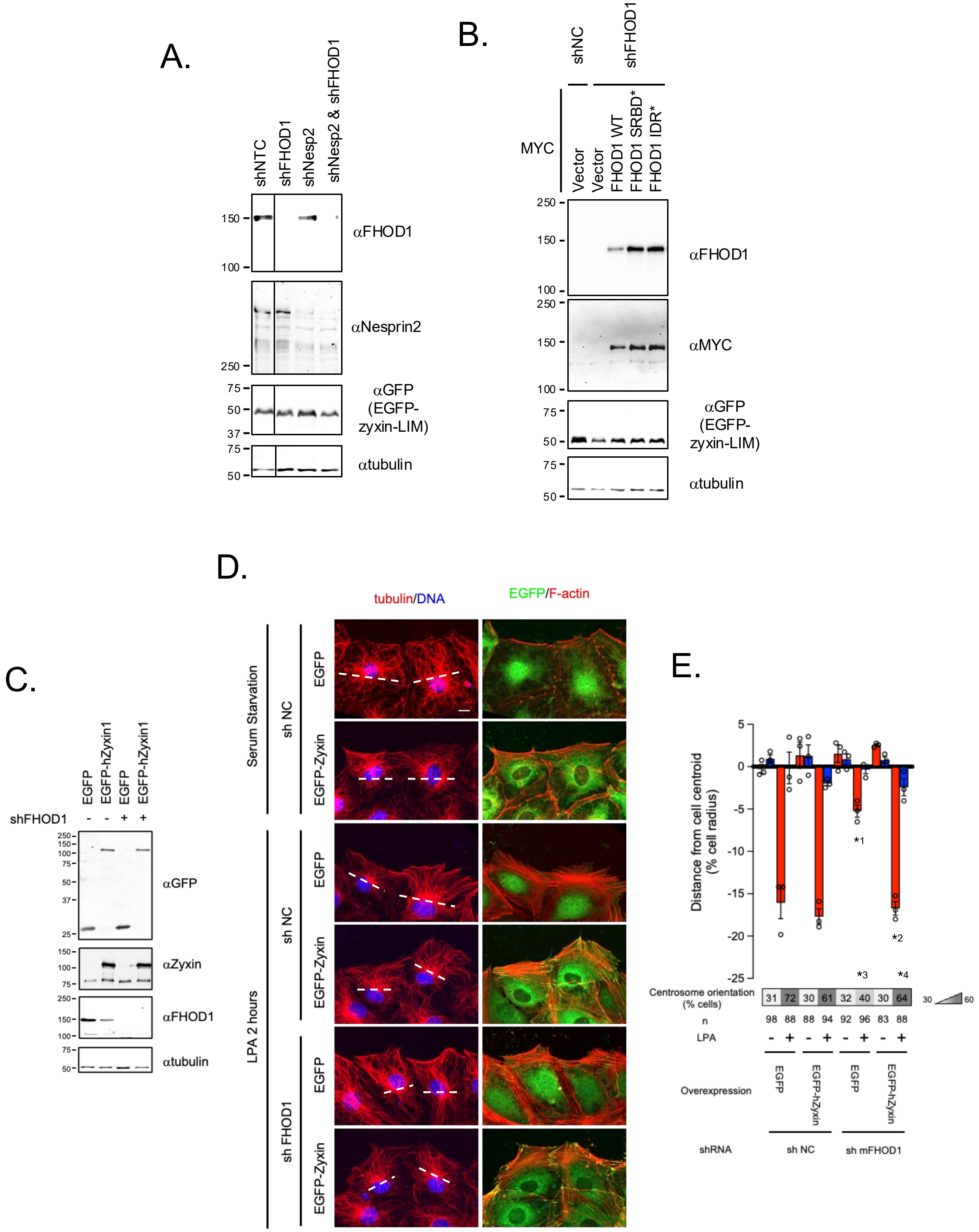

Supplemental Figure 6.

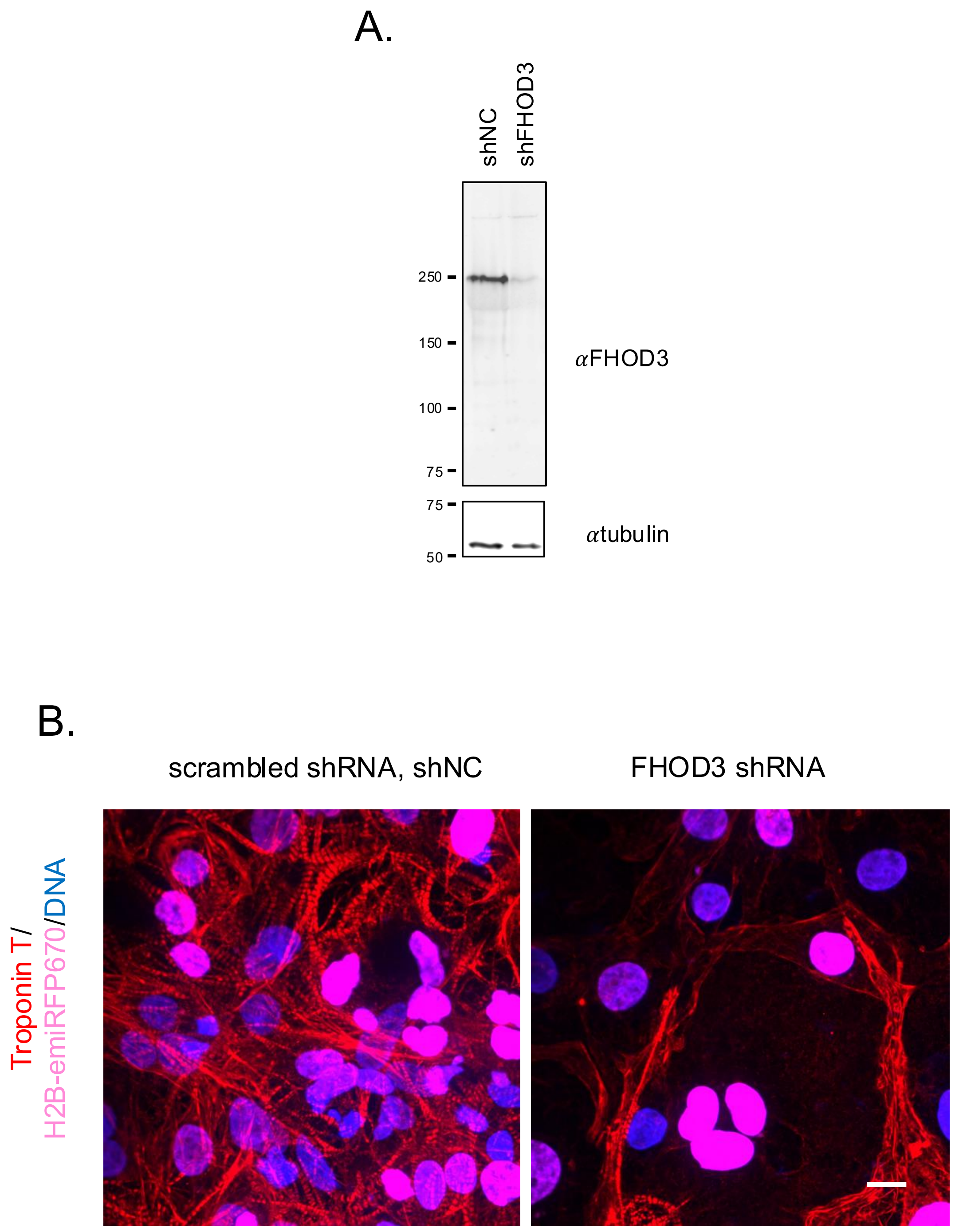

Supplemental Figure 7.

A.

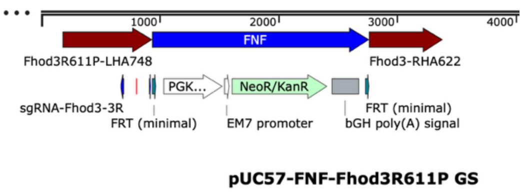


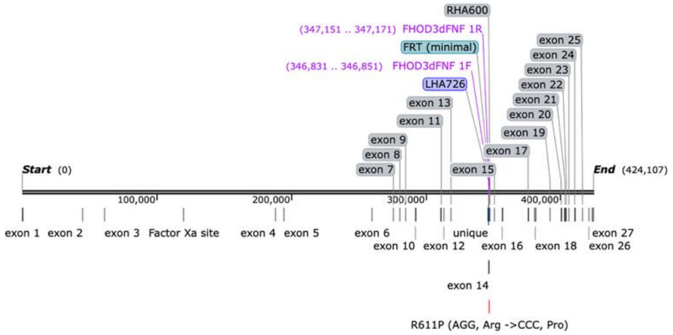


B.

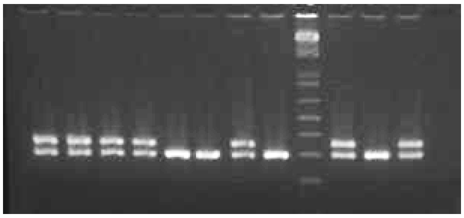

C.

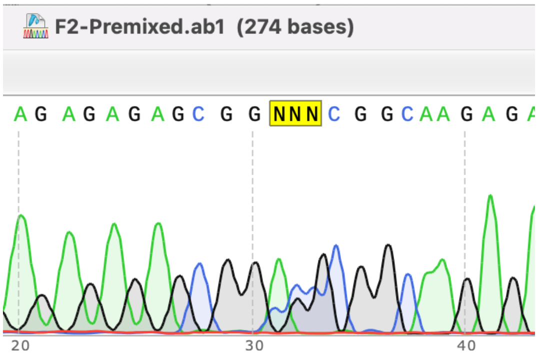

### Supplemental Figure 8.

A.

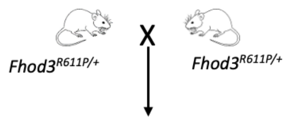


| | *Fhod3*$^{+/+}$ | *Fhod3*$^{R611P/+}$ | *Fhod3*$^{R611P/R611P}$ |
|---|---|---|---|
| Number of mice | 40 | 102 | 0.0 |
| Actual ratio | 0.28 | 0.72 | 0.0 |
| Expected ratio | 0.25 | 0.50 | 0.25 |

Chi-square= 49.61
$p < 0.0001$

B.

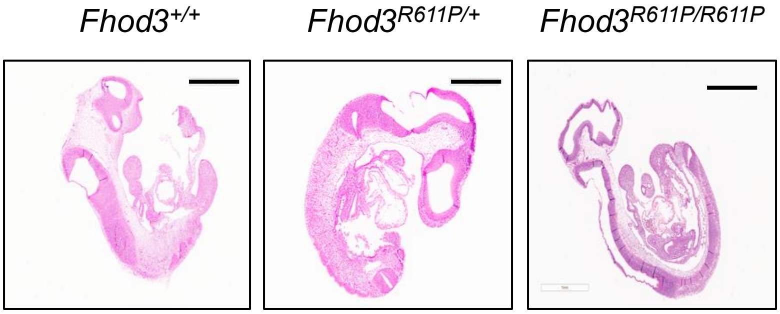


C.

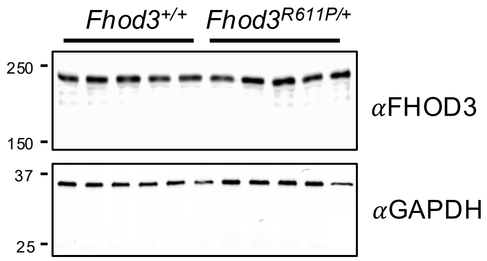


D.

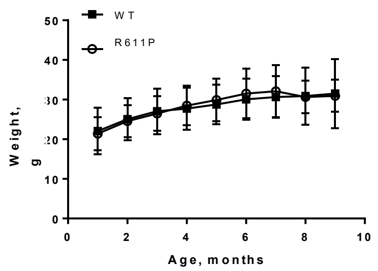


E.

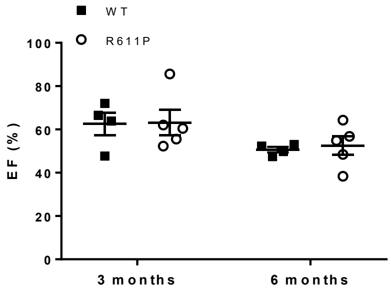


F.

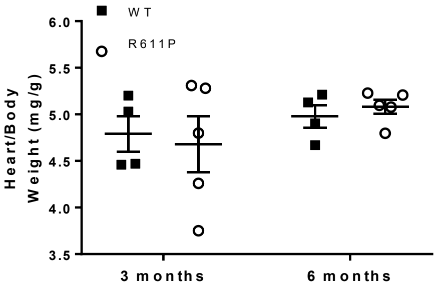


G.

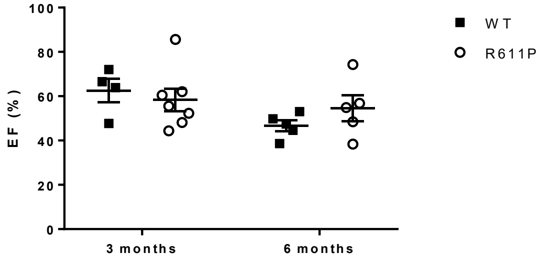

Supplemental Figure 9.

A.

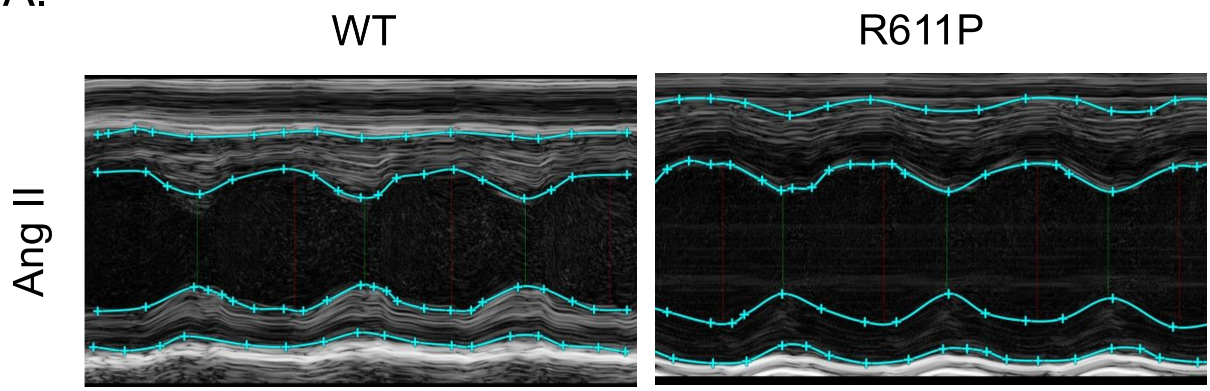


B.

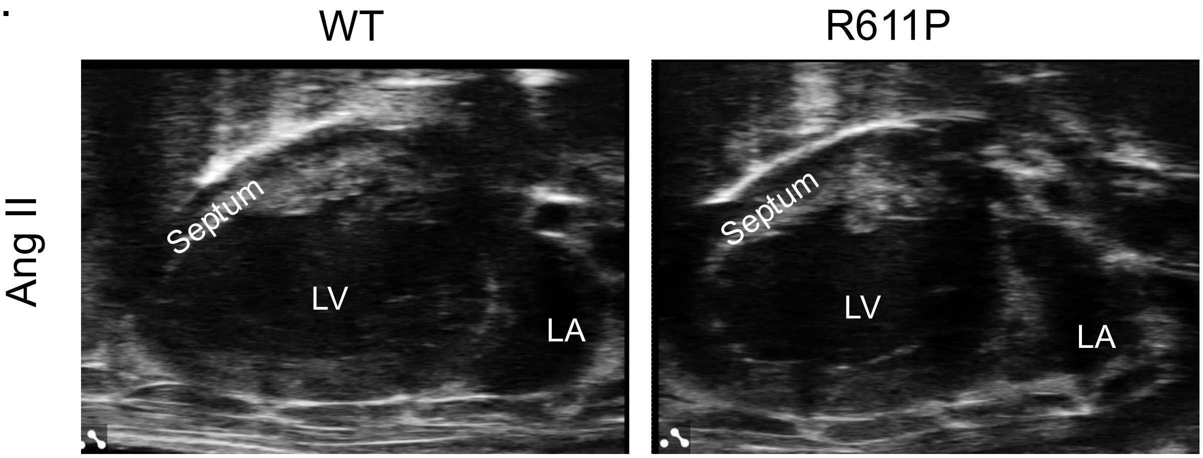
